# High-throughput mutagenesis identifies mutations and RNA-binding proteins controlling *CD19* splicing and CART-19 therapy resistance

**DOI:** 10.1101/2021.10.08.463671

**Authors:** Mariela Cortés-López, Laura Schulz, Mihaela Enculescu, Claudia Paret, Bea Spiekermann, Anke Busch, Anna Orekhova, Fridolin Kielisch, Mathieu Quesnel-Vallières, Manuel Torres-Diz, Jörg Faber, Yoseph Barash, Andrei Thomas-Tikhonenko, Kathi Zarnack, Stefan Legewie, Julian König

## Abstract

During CART-19 immunotherapy for B-cell acute lymphoblastic leukaemia (B-ALL), many patients relapse due to loss of the cognate CD19 epitope. Since epitope loss can be caused by aberrant *CD19* exon 2 processing, we herein investigate the regulatory code that controls *CD19* splicing. We combine high-throughput mutagenesis with mathematical modelling to quantitatively disentangle the effects of all mutations in the region comprising *CD19* exons 1-3. Thereupon, we identify ~200 single point mutations that alter *CD19* splicing and thus could predispose B-ALL patients to CART-19 resistance. Furthermore, we report almost 100 previously unknown splice isoforms that emerge from cryptic splice sites and likely encode non-functional CD19 proteins. We further identify *cis*-regulatory elements and *trans*-acting RNA-binding proteins that control *CD19* splicing (e.g., PTBP1 and SF3B4) and validate that loss of these factors leads to enhanced *CD19* mis-splicing. Our dataset represents a comprehensive resource for potential prognostic factors predicting success of CART-19 therapy.

**Highlights:**

- Mutations in relapsed CART-19 patients lead to *CD19* mis-splicing
- High-throughput mutagenesis uncovers ~200 single point mutations with a potential role in CART-19 therapy resistance
- Many mutations generate non-functional CD19 proteins by activating cryptic splice sites
- RNA-binding proteins such as PTBP1 are key to the expression of properly spliced, CART-19 immunotherapy-sensitive isoforms

## Introduction

B-cell acute lymphoblastic leukaemia (B-ALL) is a hematologic malignancy which causes a significant number of childhood and adult cancer deaths. In CART-19 immunotherapy, chimeric antigen receptor-armed autologous T-cells (CARTs) are engineered to target the surface antigen CD19 on B-cells by linking the single-chain variable fragment (scFv) of an anti-CD19 antibody to the intracellular signalling domain of the T-cell receptor [1]. Upon CD19 recognition, the chimeric antigen receptors activate the cytotoxic T-cells to attack the tumour cells. CART-19 therapy was recently approved for the treatment of paediatric B-ALL in the US and Europe. Unfortunately, up to 50% of children relapse under CART-19 therapy, and response rates are even worse in adults [2,3]. Several studies reported that in 40-60% of cases the cancerous B-cells get invisible to the CARTs due to loss of detectable CD19 epitope (CD19-negative) [4–7]. This recurrently involves alternative splicing of the *CD19* pre-mRNA [8–10].

Splicing comprises the excision of introns and the joining of exons by the spliceosome to generate mature mRNAs. During alternative splicing, certain exons can be either included or excluded (“skipped”), thus leading to different transcript isoforms. The splicing outcome at each exon is controlled by a large set of *cis*-regulatory elements in the RNA sequence which are recognised by *trans*-acting RNA-binding proteins (RBPs) that guide the spliceosome activity. It is increasingly recognised that widespread alterations in splicing are a molecular hallmark of cancer and often contribute to therapeutic resistance (reviewed in [11]). For instance, intron retention, i.e., the failure to remove certain introns, often disrupts the open reading frame with premature termination codons (PTCs) and thereby compromises the expression of the encoded proteins. Consistent with the widespread splicing changes, cancer-causing driver mutations frequently occur in splice-regulatory *cis*-elements, and many splicing factors have oncogenic properties, being commonly mutated or dysregulated in cancer [11–13].

Multiple alternative splicing events in *CD19* mRNA have been described to interfere with CART-19 therapy [8,10,14–17]. Most prominently, skipping of exon 2 results in a truncated CD19 protein which is no longer presented on the cell surface and hence fails to trigger CART-19-mediated killing [8,14]. In addition, it was reported that relapsed patients showed retention of intron 2 which introduces a PTC, thereby disrupting CD19 expression [10]. Similarly, simultaneous skipping of exons 5 and 6 introduces a PTC [8]. The splicing alterations can be caused by mutations within the *CD19* gene or by changes in the expression of *trans*-acting RBPs. For instance, it has been suggested that the known splicing regulator SRSF3 binds to *cis*-regulatory elements within *CD19* exon 2 to promote its inclusion [8]. Of note, alternative *CD19* isoforms showing exon 2 skipping were observed to pre-exist in patients prior to CART-19 therapy [15,16], suggesting that *CD19* splicing patterns may harbour prognostic information and could be modulated to re-establish sensitivity to CART-19 mediated killing. However, Orlando and co-workers suggested that alternative splicing changes in B-ALL patients are present in diagnostic samples at low frequency and may not contribute meaningfully to CD19 epitope loss [4]. We therefore set out to investigate *CD19* alternative splicing and its molecular determinants in B-ALL in more detail.

High-throughput mutagenesis screens combined with next-generation sequencing provide comprehensive insights into the regulatory code of splicing [18–21]. The interpretation of such data is challenging, as the mutation effects often depend on other mutations and are typically most pronounced at intermediate exon inclusion levels [18,19,22]. We and others have shown by mathematical modelling that kinetic models account for the context-dependence of mutation effects on splice isoforms [18,19]. By these models, systems-level insights can be gained into complex *cis*-regulatory landscapes, effects of *trans*-acting RBPs and principles of splicing regulation [18,19,23].

In this manuscript, we combine B-ALL patient data with high-throughput mutagenesis, mathematical modelling and RBP knockdowns to comprehensively characterise *cis*-regulatory mutations and *trans*-acting RBPs controlling *CD19* exon 2 splicing. Unlike previous mutagenesis screens, we determine all intronic and exonic mutation effects in a 1.2 kb region and quantify the abundance of 100 alternative isoforms, including intron 2 retention and alternative 3’/5’ splice site usage. Many of these isoforms encode for a non-functional CD19 protein and are therefore likely to impair CART-19 therapy. By *in silico* analyses and RBP knockdowns, we identify *trans*-regulators of *CD19* splicing that promote the production of the therapy-relevant isoforms. Taken together, our dataset is a comprehensive resource for prognostic markers of CART-19 therapy resistance and for a systems-level understanding of the splicing code.

## Results

### CART-19 patients show increased *CD19* intron 2 retention after relapse

To resolve the contribution of *CD19* splicing in CART-19 therapy, we re-analysed RNA-seq data from Orlando and co-workers [4], in which B-ALL cells of 17 patients were sequenced at initial screening and after relapse. In contrast to the original study, we expanded the analyses to intron retention surrounding *CD19* exon 2. We found that the average frequency of retention of intron 2 across patients significantly increases from 63% before therapy to 82% after relapse (*P* value = 0.022, Wilcoxon signed-rank test; **Figure 1A, B**). The trend towards higher intron 2 retention is preserved in 7 out of 10 individual patients that were sequenced both before therapy and after relapse (**Figure 1B**). Since the resulting isoform does not encode the CD19 epitope, this suggests that increased intron 2 retention contributes to CART-19 therapy relapse as reported in a recent study [10].

**Figure 1.**
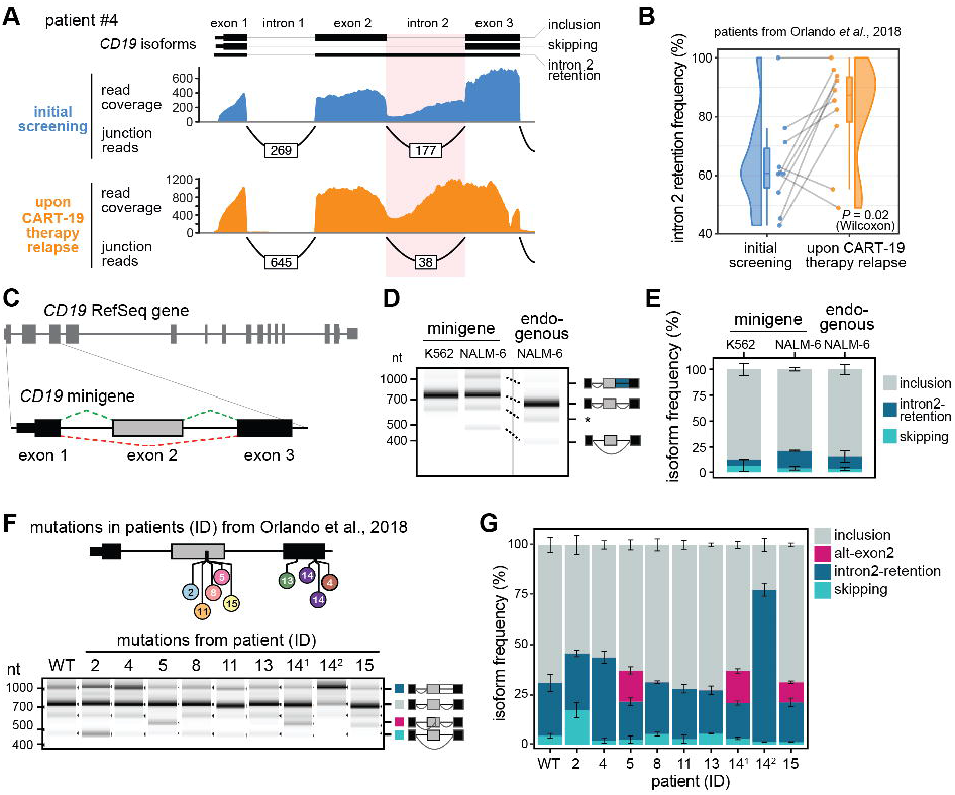
Mutations from B-ALL patients cause *CD19* mis-splicing. **(A)** Patient #4 shows increased *CD19* intron 2 retention after CART-19 therapy relapse, evidenced by reduced junction-spanning reads and increased intron coverage. Re-analysed RNA-seq data from Orlando et al. [4]. Selected isoforms (GENCODE) are shown above. **(B)** Intron 2 retention increases in B-ALL patients after CART-19 therapy relapse. Intron 2 retention frequency (as % of all isoforms) is shown for 10 patients with matched RNA-seq data at screening and after relapse. *P* value = 0.02, paired Wilcoxon signed-rank test. **(C)** The *CD19* minigene spans exons 1-3 and the intervening introns from the *CD19* gene. **(D, E)** The minigene generates the same isoforms as the endogenous *CD19* gene in NALM-6 cells. Gel-like representation (D) and quantification (E) of semi-quantitative RT-PCR showing detected isoforms intron2-retention (blue), inclusion (grey) and skipping (turquoise) for the WT minigene in NALM-6 and K562 cells as control. Isoforms of *CD19* gene in NALM-6 cells are shown for comparison. Asterisk indicates a previously reported RT-PCR artefact [57] (see methods). Error bars indicate standard deviation of mean (s.d.m.), n = 3 replicates. **(F, G)** Patient mutations cause splicing changes in the *CD19* minigene. Top: Location of the tested mutations. Numbers refer to patient IDs as reported in Orlando et al. [4]. 14.1 and 14.2 correspond to distinct mutations from patient #14. Gel-like representation (F) and quantification (G) of semi-quantitative RT-PCR as in (D) and (E). Additional isoform alt-exon2 (purple) includes a truncated version of exon 2. Error bars indicate s.d.m., n = 3 replicates.

### Somatic mutations in relapsed patients cause splicing alterations

The majority of relapsed patients in the Orlando study (12 out of 17) [4] harbour somatic mutations within the *CD19* gene, including frameshift insertions, deletions and single nucleotide missense variants. We selected nine mutations in exons 2 or 3 from eight patients for further analysis (**Table S1**). To test for effects on splicing, we constructed a minigene reporter that harbours *CD19* exon 1-3 including the two intervening introns 1 and 2 (**Figure 1C**). We confirmed that the minigene gives rise to the same transcript isoforms as the endogenous gene in the human B-ALL cell line NALM-6 (**Figure 1D, E**). When introducing the patient mutations into our minigene reporter, we found that six out of nine tested mutations lead to the production of alternative *CD19* isoforms linked to CART-19 therapy resistance (**Figure 1F, G**): The mutation from patient #2 induces exon 2 skipping, while mutations from patients #4 and #14.2 cause intron 2 retention. In addition, three mutations enhance the production of an additional isoform that uses an alternative 3’ splice site in exon 2 (termed alt-exon2; mutations from patients #5, #14.1 and #15). The alternative splice junction in alt-exon2 introduces a frameshift causing a PTC and will hence abolish the production of a targetable CD19 epitope. We note that as reported by Orlando and co-workers [4], most of the tested mutations also introduce frameshifts, making it difficult to discriminate between PTC-induced and splicing-mediated defects. For instance, the alternative 3’ splice site of alt-exon2, which is prevalent in patient #5, in fact compensates for the frameshift that is introduced by the concomitant deletion, i.e., restores the open reading frame (**Figure S1A**). Thus, taking the splicing information into account changes the interpretation of what CD19 protein variants are expressed in a given patient. More broadly speaking, these results suggest that *CD19* mutations in CART-19 relapse patients frequently trigger splicing changes that potentially influence therapeutic outcomes.

### High-throughput screening of *CD19* exons 1-3 alternative splicing

To systematically study the effects of point mutations on *CD19* exons 1-3 splicing, we adopted our previously developed massively parallel splicing reporter assay [18] (**Figure 2A**). To this end, we randomly introduced point mutations as well as short insertions and deletions into the *CD19* minigene reporter by error-prone PCR. This yielded a pool of 10,295 minigene variants, each with a different set of mutations and tagged with a unique 15-nt barcode sequence. As an internal control, 194 wild type (WT) minigenes with distinct barcodes were added. Mutations in all minigene variants were mapped using targeted long-read DNA sequencing (DNA-seq, PacBio SMRT-seq, **Figure S1B, C**) and validated for 30 minigene clones via Sanger sequencing. The DNA-seq data shows that the minigene variants contain on average 9.7 mutations (**Figure S1D**). This allows for a comprehensive characterisation of the mutation landscape, as each position is on average mutated in 80 different minigene variants and 90% of the mutations are present in at least four distinct minigene variants (**Figure S1E, F**). To measure splicing outcomes, the minigene pool was transfected into NALM-6 cells and the resulting transcripts were quantified by targeted RNA sequencing (RNA-seq) using 350 nt + 250 nt paired-end reads (Illumina MiSeq, **Figure S1B, S2A**). We detected around 100 different splice isoforms (see below) which were unambiguously identified by paired-end sequencing. Two replicate experiments showed high correlation in the measured isoform frequencies (R between 0.91 and 0.98 for the different isoforms, **Figure S2B**). Based on the common barcode sequence, information from DNA and RNA sequencing could be combined, linking mutations at the DNA level to frequencies of RNA splice isoforms for a total of 10,295 minigenes in two replicate experiments (**Table S2**).

**Figure 2.**
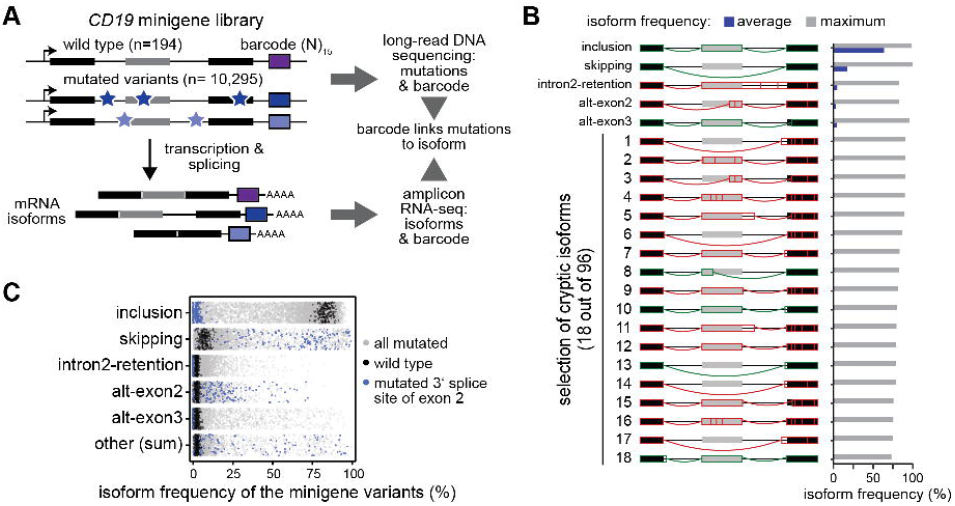
High-throughput mutagenesis identifies splicing-effective mutations and cryptic isoforms in the *CD19* minigene. **(A)** High-throughput detection of splicing-effective mutations and cryptic isoforms. Mutagenic PCR creates mutated minigene variants (top) that upon transfection into NALM-6 cells give rise to alternatively spliced transcripts (bottom). Mutations (stars) and corresponding splicing products are characterised by DNA and RNA sequencing, respectively, and linked by a unique 15-nt barcode sequence in each minigene (coloured boxes). Black and grey boxes depict constitutive and alternative exons, respectively. **(B)** A large number of *CD19* splice isoforms arise in the minigene library. *CD19* splice isoforms with highest maximal isoform frequency across all 9,321 minigene variants. Schematic representation (left) of 5 major and 18 cryptic isoforms depicts exons 1-3 (boxes) and introns (horizontal lines) with splice junctions for each isoform (arches). Colour indicates coding potential (green, coding; red, non-coding). Bar graph (right) shows average and maximal isoform frequency across all minigenes. Cryptic isoforms are sorted by maximal isoform frequency (**Table S3**). **(C)** Inclusion isoform dominates in WT minigenes, whereas mutated variants show broad spread in all major isoforms. Frequencies of five major isoforms in replicate 1 for all wild type (black; n = 195) and mutated (grey; n = 9,476) minigenes in the library. Minigene variants harbouring a mutation in the 3’ splice site of exon 2 (n = 174) are highlighted in blue. “Other” refers to the sum of 96 cryptic isoforms.

### Therapy-relevant isoforms accumulate in response to numerous point mutations

To our surprise, the screen revealed a high complexity of *CD19* exon 1-3 splicing, with a total of 101 alternative isoforms occurring with a frequency of ≥5% of all transcripts in at least two minigene variants (**Table S3**). Out of these, the five major isoforms exceed 1% in WT minigenes, whereas the others, termed cryptic isoforms, only accumulate in mutated minigene variants (**Figure 2B**). In WT, the by far most abundant major isoform is exon 2 inclusion (termed “inclusion”, followed by exon 2 skipping (termed “skipping”) and intron 2 retention (termed “intron2-retention”). Two additional major isoforms in WT originate from alternative 3’ splice site usage within exon 2 (alt-exon2) and 3 (alt-exon3) (**Figure 2B, C**). Notably, alt-exon2 is the same isoform as observed upon patient mutations above. As expected, the measured frequencies for the major isoforms show little variance for the 194 unmutated WT minigenes (standard deviation < 6%, **Figure 2C**). In contrast, many mutated minigene variants show strong changes relative to WT, suggesting a large impact of specific mutations on splicing outcomes (**Figure 2C**). For instance, all minigenes with a mutation in the 3’ splice site of exon 2 lose the inclusion isoform, accompanied by strong alterations in the remaining major isoforms. Taken together, these observations support the accuracy of our screening results.

All major isoforms, except exon 2 inclusion, encode for a truncated CD19 receptor lacking a functional CART-19 epitope and could thus contribute to therapy resistance. Our unbiased screening approach extends the list of potentially therapy-relevant *CD19* mutations, since 1,721 out of 9,127 mutated minigenes show exon 2 skipping, intron 2 retention and/or alt-exon2 isoform frequencies of >25% (**Figure 2C**). However, since the minigene variants carry on average 9.7 point mutations, the observed splicing changes represent the combined effects of several mutations. To extract the impact of individual mutations, we adapted our previous mathematical modelling framework [18] and implemented a multinomial logistic regression approach. Here, the splicing change in each minigene variant is described as the sum of the underlying point mutation effects (**Figure 3A**, see Methods). These single mutation effects are unknown and are determined by simultaneously fitting the model to all minigene measurements. Thereby, we were able to infer the individual effects of 4,255 point mutations on the five major isoforms (**Figure 3A, S3A**). We validated the reliability of this model in describing combined mutations using a 10-fold cross-validation approach, in which we left out 10% of all minigene variants from fitting and were able to accurately predict them after model fitting (Pearson correlation coefficients 0.65-0.95; **Figure 3B, S3B**). Furthermore, the model performed well in predicting single mutation effects, as soon as a mutation occurred in three or more minigenes in the dataset (**Figure S4C**), which applied to 90% of all mutations (**Figure S1F**).

**Figure 3.**
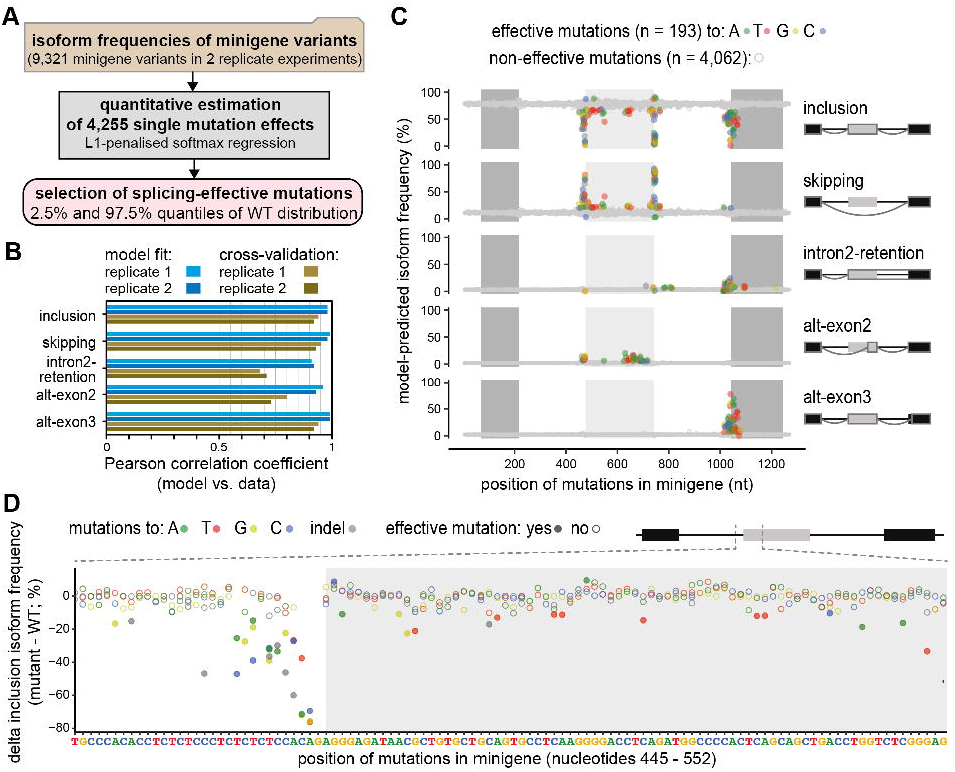
Quantitative modelling predicts single mutation effects on splice isoforms. **(A)** Multinomial logistic regression workflow for the quantification and selection of single mutation effects. Based on the experimentally measured frequencies of five major isoforms in 9,321 minigene variants (top box), a softmax regression model was formulated to estimate 4,255 single mutation effects from the data (middle box) using L1 penalisation to prevent overfitting. Splicing-effective mutations were selected for each isoform based on the respective empirical WT frequency distribution using the 2.5% and 97.5% quantiles as cutoff. **(B)** Splicing-effective mutations accumulate in distinct regions around exons 2 and 3. Landscape of model-predicted single mutation effects on five major isoforms (indicated on the right). Predicted isoform frequencies are plotted as a function of the position of a mutation. Colours indicate the nucleotide substitution of splicing-effective point mutations (see legend), whereas non-effective mutations are grey. **(C)** The model performs well in fitting and 10-fold cross-validation. Bars show Pearson correlation coefficients between model and data for two replicates and each of the five isoforms across all combined mutation minigenes considered in model training and validation, respectively. See **Figure S3A, B** for corresponding scatter plots. **(D)** Zoom-in shows the model-predicted delta inclusion isoform frequency (frequency for a point mutation - frequency in WT) for nucleotides 445-552 of the minigene. The type of nucleotide substitution is shown for all mutations, with splicing-effective mutations highlighted as filled circles.

Out of 4,255 quantified single mutation effects, we find 193 splicing-effective mutations that significantly alter the frequency of at least one isoform in the two replicates beyond the 2.5 and 97.5% quantiles of the WT minigene distribution (**Figure 3C, Table S4, Data S1**). 33 of these splicing-effective mutations overlap with single nucleotide variants (SNVs) that were previously reported in the human population from whole-genome or exome sequencing data (**Table S5**). The strongest mutation effects accumulate around the four main splice sites and throughout exon 2 and correspond to the core *cis*-regulatory elements, such as splice-site dinucleotides, branchpoint and polypyrimidine tract, as well as auxiliary elements (**Figure 3C, D**). Inspecting in more detail the 83 mutations that specifically impact on *CD19* exon 2 skipping, we find them to cluster within and around exon 2. In particular, 21% of all positions within exon 2 (55 out of 267 nt) harbour at least one splicing-effective mutation, suggesting that *CD19* exon 2 is densely packed with *cis*-regulatory elements controlling its inclusion. In addition, we observe smaller clusters of mutations within the introns and flanking constitutive exons which likely represent more distal *cis*-regulatory elements (**Figure 3C**). Similarly, we explored the 54 splicing-effective mutations that impact on intron 2 retention. As expected, strongest effects are observed at the splice sites of intron 2. In addition, we find clusters of mutations in intron 2 and exon 3 that might reflect important *cis*-regulatory elements. The effect of all mutations on the five major isoforms can be explored in **Data S1**.

In conclusion, our combined screening and modelling approach quantitatively describes alternative splicing of *CD19* exons 1-3 by predicting the effects of all individual point mutations and combinations thereof. Our screen thereby represents a comprehensive resource for the identification of mutations with clinical relevance in CART-19 therapy resistance.

### Cryptic isoforms destroy the *CD19* ORF and are associated with recurrent mutations

Besides the five major isoforms, the *CD19* exons 1-3 can give rise to 96 cryptic isoforms which are rare (<1%) in WT, but accumulate upon certain mutations (**Figure 2B, Table S3**). The cryptic isoforms involve a total of 71 cryptic splice sites (**Figure 4A**). Of note, 33 of these cryptic isoforms make up more than 50% of total transcripts and are therefore dominant in certain minigene variants (**Figure 2B, C**). To assess whether these cryptic isoforms impact on CD19 epitope presentation, we analysed their coding potential and found that the vast majority of cryptic *CD19* isoforms (78 out of 96) show a frameshift and/or carry a PTC (**Figure 4B**). This will either lead to the production of truncated CD19 peptides that likely do not allow for presentation on the cell surface [14] or will induce nonsense-mediated mRNA decay of the cryptic isoforms and will hence reduce *CD19* transcript and protein levels.

**Figure 4.**
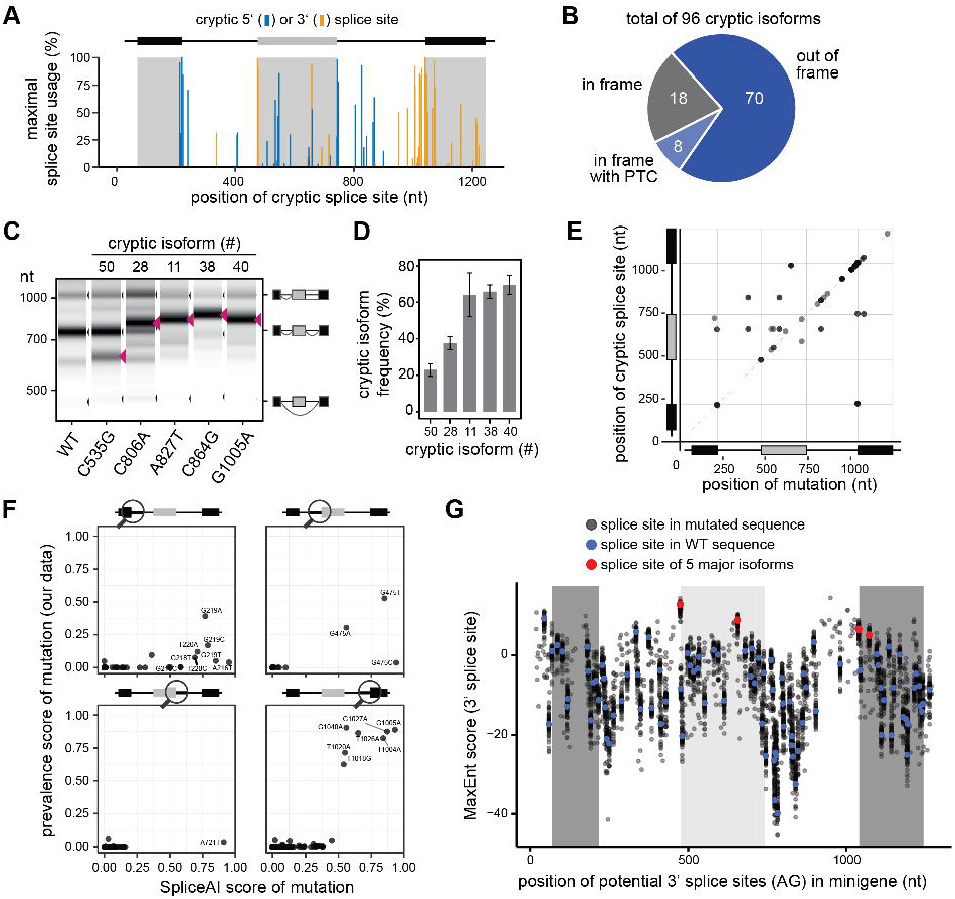
*CD19* mutations frequently activate cryptic splice sites. **(A)** Alternative splicing of *CD19* minigene variants involves 71 cryptic splice sites. Splice site usage was calculated for each minigene variant by dividing the sum of junction reads involving a particular splice site by the total number of reads. The maximum usage across all minigenes is plotted against the corresponding position to the cryptic splice sites. **(B)** Cryptic isoforms code for non-functional CD19 proteins. Out of 96 cryptic isoforms, 8 run into a premature termination codon (PTC) and 70 are out-of-frame, thus potentially encoding non-functional CD19 protein variants. The remaining 18 remain in frame, but are shortened or extended relative to the reference inclusion isoform. **(C, D)** Experimental validation of five point mutations that are associated with distinct cryptic isoforms. Targeted point mutations were introduced into the *CD19* minigene, and splicing outcomes were determined by semi-quantitative RT-PCR. Predicted cryptic isoforms are indicated by red arrowheads. Gel-like representation (C), with major isoforms indicated on the right, and quantification (D). Error bars indicate s.d.m., n = 3 replicates. **(E)** Mutations leading to cryptic isoforms are often located within or near cryptic splice sites. For 31 cryptic isoforms that are highly associated with a mutation (prevalence score > 0.25; y-axis), the position of this mutation (x-axis) was related to the position of the used cryptic splice site (y-axis). **(F)** SpliceAI correctly predicts single mutations leading to the generation of cryptic isoforms. SpliceAI was used to predict changes in splice junctions based on pre-mRNA sequence for all possible *CD19* minigene single mutants. SpliceAI scores of 0 and 1 reflect 0% or 100% probability to gain a cryptic splice site in response to a mutation, respectively (see Methods). Scatter plots compare the SpliceAI score against the prevalence score from our data, which quantifies the association of a mutation with a cryptic isoform, for 254 mutation-splice site pairs that match in their positions with SpliceAI. Separate panels are shown for each region around a canonical splice site (circle in schematic minigene representation). **(G)** Exon 3 harbours a weak 3’ splice site and is preceded by a high number of potentially competing cryptic 3’ splice sites, which often reach similar strength upon mutation. Dotplot shows splice site strengths (MaxEnt score) for putative 3’ splice sites (AG dinucleotides) in the *CD19* minigenes. MaxEnt score was calculated in a 23-nt sliding window for the WT sequence (red and blue dots) and hypothetical mutant minigenes, in which all possible single point mutations were introduced (grey dots). The 3’ splice sites used in the five major isoforms are highlighted in red.

To derive a mechanistic understanding of cryptic isoform biogenesis, we analysed the underlying point mutations. To this end, we calculated a prevalence score which quantifies the degree of association between an isoform and a point mutation. This was done based on the measured isoform frequencies in the minigene library by multiplying: (i) the frequency of a mutation being present if the isoform level is high (>5%), and (ii) the frequency of the isoform level being high given that the mutation is present. A prevalence score of 1 indicates perfect correspondence between mutation and isoform, whereas a prevalence score of 0 is observed if they are unrelated. This score-based analysis showed that 36 cryptic isoforms are specifically associated with 31 specific point mutations (38 mutation-isoform pairs with prevalence score > 0.25, **Figure S4A, Table S3**). The remaining 60 cryptic isoforms do not show a specific association, implying that they can either be generated by multiple redundant mutations, or that our screen lacks sufficient coverage to support a reliable association. To directly test the predicted associations, we introduced five mutations with a specific association to a cryptic isoform in our minigene reporter (C535G, chr16:28932405, prevalence score = 0.18; C806A, chr16:28932676, 0.68; A827T, chr16:28932697, 0.93; C864G, chr16:28932875, 1; G1005A, chr16:28932734, 0.89). Semi-quantitative RT-PCR confirmed that all five tested mutations lead to the appearance of the associated cryptic isoform (**Figure 4C, D**).

Altogether, our analysis provides a list of 31 mutations that are likely to trigger cryptic isoform formation. Importantly, the resulting cryptic isoforms show a maximum usage of up to 91% (**Table S3**), which is likely to drastically interfere with normal *CD19* splicing, protein production and subsequent epitope presentation. The associated mutations may thus provide predictive biomarkers for CART-19 therapy response in the future.

### The cryptic isoforms are caused by mutations that disrupt or create splice sites

Due to their potential clinical relevance, we wanted to learn more about how the mutations activate the cryptic isoforms. We found that the majority of mutations with a prevalence score > 0.25 are either in close proximity or directly overlap with the associated cryptic splice site (78.9% with distance < 5 nt; **Figure 4E**). Further inspection showed that the underlying mutations either destroy the original splice site (7.9%) or generate a new cryptic splice site (57.9%). Hence, the cryptic isoforms do originate from the generation or destruction of core *cis*-regulatory elements rather than affecting auxiliary elements.

Currently, major efforts are ongoing to implement artificial intelligence (AI) tools to predict the effect of clinical variants on the splicing outcome. We therefore tested whether the state-of-the-art neural network [24], which predicts changes in the splicing patterns induced by single point mutations, captures the gain and loss of splice sites in *CD19*. To this end, we applied SpliceAI using all possible single point mutations in the *CD19* minigene as an input. Similar to the results from our mutagenesis screen (**Figure 4A**), SpliceAI predicts cryptic splice site activation by mutations throughout the minigene, with an increased density around the 3’ splice site of exon 3 (**Figure S4B**). All SpliceAI-predicted mutations are close to the affected cryptic splice sites (**Figure S4C**). Hence, SpliceAI successfully reflects the global landscape of mutation-induced cryptic splice site activation in the *CD19* minigene.

With respect to the accuracy of the individual predictions, we found that 10 out of 38 mutations with strong SpliceAI predictions (SpliceAI score > 0.5) indeed lead to the accumulation of splice isoforms with the corresponding cryptic splice sites in the experimental data (prevalence score > 0.25, **Figure 4F**). In the remaining 28 cases, either weak overall cryptic splice site activation occurred in the data (9 cases) or a different cryptic splice site was activated than predicted by SpliceAI (19 cases; **Figure S4B**). In quantitative terms, the likelihood of a cryptic splice site activation according to the SpliceAI prediction (“SpliceAI score”) is correlated to the magnitude of the prevalence score linking the mutation to the corresponding cryptic isoform in our screen (**Figure 4F**). Overall, the comparison supports that SpliceAI can guide the interpretation of mutation effects in clinical samples, though direct experimental validation is necessary. As such, our data can be used to benchmark new tools for splicing prediction.

The cryptic isoforms arise from numerous 3’ and 5’ cryptic splice sites that distribute over the entire minigene and accumulate at exon 3 (**Figure 4A**). In line with a high penetrance, 26 cryptic splice sites reach more than 50% usage upon certain mutations, particularly around the start of exon 3. We hypothesised that cryptic splice site activation occurs in exon 3 because its canonical splice site can be outcompeted by neighbouring cryptic sites. To test this, we scored the strength of local consensus sequences using MaxEntScan [25], and indeed found that the 3’ splice site of exon 3 is weak compared to all other canonical splice sites of *CD19* exons 1-3 (**Figure 4G, S4D, E**). In line with our hypothesis, mutations around the 3’ splice site of exon 3 frequently create stronger splice sites than elsewhere in the minigene that exceed the strength of the canonical 3’ splice site of exon 3 (**Figure 4G**). This suggests that weak splice sites are particularly vulnerable for the activation of competing cryptic splice sites and should be of particular interest when assessing the impact of clinical variants on splicing outcomes.

### An extensive network of RBP regulators might drive *CD19* mis-splicing

Besides *CD19* mutations, CART-19 therapy resistance may also stem from altered expression of *trans*-acting RBPs which bind to the *CD19* pre-mRNA to control alternative splicing. To identify putative RBP regulators, we explored publicly available databases containing experimentally determined RBP binding motifs (ATtRACT [26], oRNAment [27]). Furthermore, we employed DeepRiPe [28], a neural network-based algorithm trained on PAR-CLIP and ENCODE eCLIP datasets that predicts changes of RBP binding upon mutation. In combination, these tools predict a total of 198 RBPs to bind within *CD19* exons 1-3 (ATtRACT: 62 RBPs; oRNAment: 70 RBPs) or to change binding upon mutation (DeepRiPe: 128 RBPs; **Figure 5A-C, S5A**).

**Figure 5.**
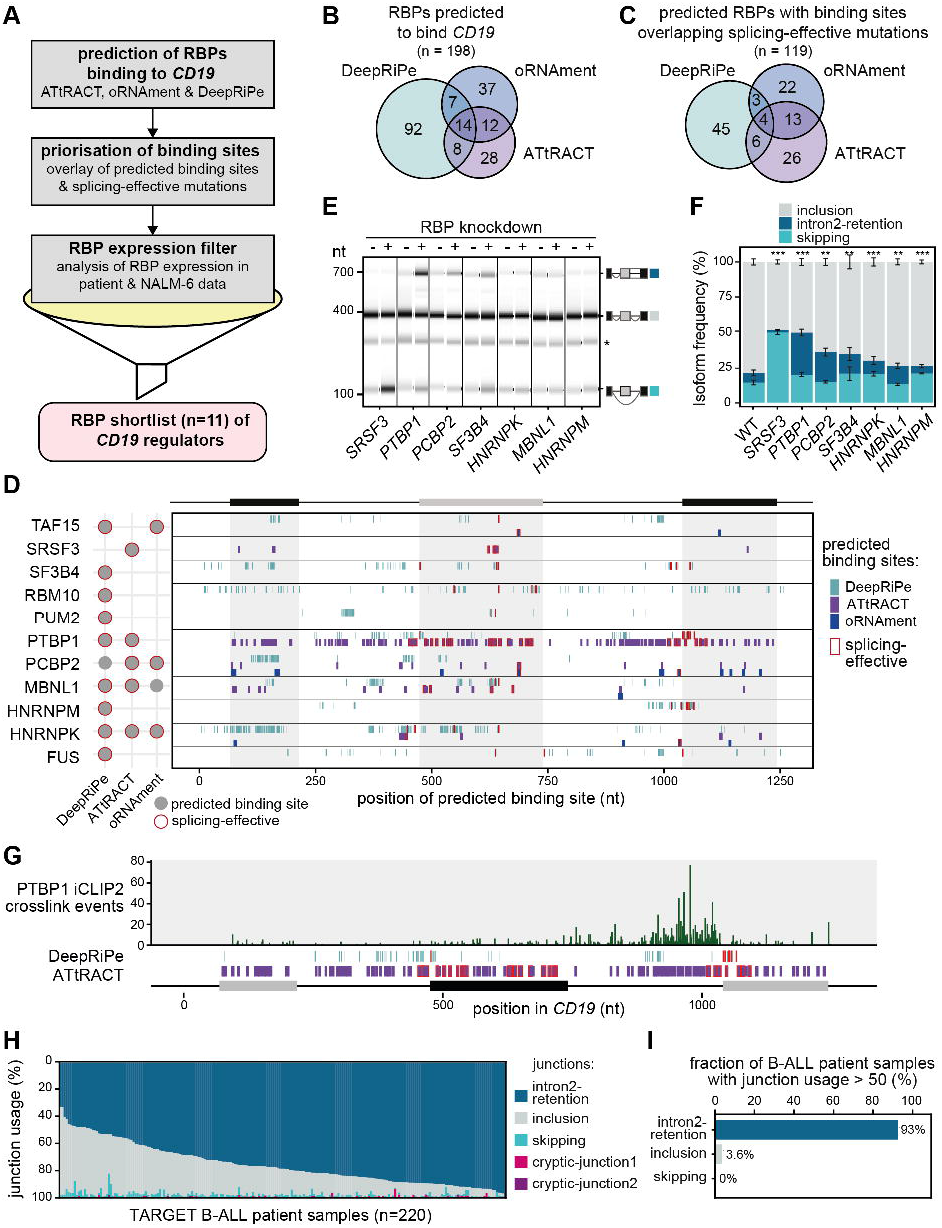
*In silico* predictions identify RBP regulators of *CD19* alternative splicing. **(A)** Pipeline for the identification of potential RBP regulators of *CD19* splicing. Starting with *in silico* predictions, we obtained 198 candidate RBPs with predicted binding motifs (ATtRACT/oRNAment) or predicted differential binding upon mutation (DeepRiPe). These were prioritised by overlapping with the splicing-effective mutations from our screen. Additionally, based on publicly available RNA-seq data, we required a minimum mean expression in RNA-seq data from B-ALL patients [29] and NALM-6 cells [66]. Together with literature information, we shortlisted 11 candidate RBPs for knockdown (KD) experiments, including SRSF3 as a positive control. **(B, C)** *In silico* analyses predict dozens of RBPs binding to *CD19*. Venn diagrams show overlap of RBPs in initial predictions (B) and after overlay with splicing-effective mutations (C). **(D)** The 11 candidate RBPs are predicted to bind throughout the *CD19* minigene region. For each RBP, the binding sites predicted by ATtRACT and oRNAment and disrupting mutations predicted by DeepRiPe, are indicated (see legend). Sites overlapping with splicing-effective mutations are framed in red. The schematic summary (left) shows that all 11 candidate RBPs have at least one predicted site that overlaps with a splicing-effective mutation. A full list of predicted binding sites (ATtRACT/oRNAment) and differential binding mutations (DeepRiPe) is provided in **Table S6**. **(E, F)** Seven RBP KDs significantly change *CD19* splicing. Gel-like representation (E) and quantification (F) of semi-quantitative RT-PCR showing detected isoforms exon 2 inclusion (grey), intron 2 retention (blue) and skipping (turquoise) from the endogenous *CD19* gene in KD and control NALM-6 cells. Asterisk indicates a previously reported RT-PCR artefact [57] (see methods). Error bars indicate s.d.m., n = 3 replicates. ** *P* value < 0.01, *** *P* value < 0.001, Student’s *t*-test. Measurements for all 11 KD experiments are shown in **Figure S6B, C**. **(G)** PTBP1 shows extensive binding to *CD19* intron 2. Bar diagram shows the number of PTBP1 iCLIP crosslink events from NALM-6 cells on each nucleotide in endogenous *CD19* exons 1-3. Predicted PTBP1 binding motifs (ATtRACT) and mutations predicted to alter PTBP1 binding (DeepRiPe) are shown below (see legend in panel D). Nucleotide positions are given relative to minigene sequence. **(H, I)** Intron 2 retention is the predominant isoform in B-ALL patients. (H) Stacked barchart shows the relative usage (percent selected index, PSI) of all junctions originating from exon 3 (**Figure S6D**) in 220 B-ALL patients (TARGET program). (I) Barchart quantifies the fraction of patients in which a given junction rises to PSI > 50%.

To link the putative RBP regulators to the observed splicing changes, we overlaid the predicted binding sites (or predicted mutations for DeepRiPe) with splicing-effective mutations from our screen. Overall, we find that 79% and 60% of ATtRACT and oRNAment binding sites, respectively, overlap with a splicing-effective mutation (affecting any of the five major isoforms). Furthermore, 105 (5%) of the mutations predicted to change RBP binding by DeepRiPe overlap with splicing-effective mutations, suggesting that modulating RBP binding at these sites may have a functional impact on *CD19* splicing (**Figure 5A, S5A**). By merging these sets, we obtained a list of 119 RBPs that may regulate splicing by binding to *CD19* exons 1-3 (**Table S6**). Most of these are expressed in cancerous B-cells from B-ALL patients from [29] (80 with mean FPKM [fragments per kilobase of transcript per million mapped reads] > 10; **Figure S5B**) and could thus interfere with CART-19 therapy. Among these RBPs are SRSF3, a previously reported regulator of *CD19* splicing [8], but also new candidates such as PTBP1. Altogether, the *in silico* predictions suggest the presence of an extensive RBP network controlling *CD19* splicing that may impact on the CART-19 therapy success.

### Depletion of PTBP1 and several other RBPs results in non-functional *CD19* isoforms

Based on our experimental data, *in silico* predictions, expression, literature information and manual curation, we shortlisted 11 RBP candidates for further analysis, including *SRSF3* as a positive control. To test their impact on endogenous *CD19* splicing, we generated NALM-6 cell lines stably expressing shRNAs against the shortlisted RBPs (depletion to <40% transcripts; **Figure S6A**). As previously described [8], knockdown of *SRSF3* leads to increased exon 2 skipping in the endogenous *CD19* transcripts, confirming that this SR protein is required for exon 2 inclusion (**Figure 5E, F**). Importantly, we find that knockdown of six additional RBPs (PTBP1, PCBP2, SF3B4, HNRNPK, MBNL1 and HNRNPM) has significant effects on *CD19* alternative splicing (**Figure 5E, F, S6B, C**). The knockdown of these factors reduces *CD19* exon 2 inclusion, while promoting intron 2 retention and/or exon 2 skipping, thus shifting the cells towards expression of relapse-associated *CD19* isoforms. This implies that reduced levels of these factors can impair targetable CD19 epitope expression.

PTBP1 stands out among the putative regulators as it shows the strongest effects on intron 2 retention, which emerged as the most prominent *CD19* mis-splicing isoform in our re-analysis of B-ALL patient data (**Figure 1B**). PTBP1 recognises clusters of UC-rich motifs [30,31]. Remarkably, ATtRACT predicts almost 100 such PTBP1 binding motifs across the studied *CD19* region, including 25 that overlap with splicing-effective mutations (**Figure 5D, Table S6**). Moreover, DeepRiPe predicts 78 mutations in 63 positions that change PTBP1 binding, out of which 10 are splicing-effective in our screen. The high number of predicted binding sites suggests a partial redundancy, indicating that PTBP1 regulation might be difficult to disrupt with individual point mutations as introduced in our screen. To experimentally test if PTBP1 binds to the predicted sites, we performed PTBP1 iCLIP2 experiments in NALM-6 cells. In line with a role in intron 2 retention, we find extensive PTBP1 binding particularly in intron 2, where it spreads over an extended cluster of predicted binding sites (**Figure 5G**). The broad binding at splicing-effective positions and beyond supports that PTBP1 is a direct and central regulator of *CD19* alternative splicing, with most prominent effects on intron 2 retention.

Given these results, we reasoned that accumulation of the *CD19* intron 2 retention isoform in B-ALL patients due to RBP dysregulation or *CD19* sequence mutations could serve as a predictive biomarker for a poor response to CART-10 therapy. To support this hypothesis, we extended our analysis of patient RNA-seq data (**Figure 1B**) to the complete panel of 220 BALL patients from the Therapeutically Applicable Research To Generate Effective Treatments (TARGET) program. Although these patients had not been treated with CART-19 yet, intron 2 retention appeared as the predominant isoform in almost all of them (**Figure 5H, I, S6D**). This supports previous findings [10,15] that unproductive *CD19* splicing disrupts CD19 epitope presentation B-ALL patients already prior to CART-19 therapy exposure. Therefore, the splicing-effective mutations and RBP regulators identified in this work may harbour prognostic information for CART-19 therapy success.

## Discussion

Massively parallel reporter assays such as our high-throughput mutagenesis screen provide comprehensive insights into the regulatory code of splicing, as they characterise the complete set of *cis*-acting sequence mutations and reveal the binding sites of *trans*-acting RNA-binding proteins (e.g., [18–20,32–34]). The interpretation of these datasets is challenging due to nonlinear interactions of individual mutation effects. For instance, competition effects in splicing reduce the impact of individual mutations at low and high isoform frequencies, i.e., depending on the mutational background [18,19]. In addition, other factors such as RBP expression patterns and cell type/tissue identity determine the effects of sequence mutations. Using kinetic modelling, we and others derived regression models taking competition in splicing into account, thereby showing that the effects of complex mutation combinations can be quantitatively described as the sum of individual mutation effects [18,19]. Thus, mutations seem to control splicing additively rather than synergistically, and this principle also holds for *CD19* splicing.

In our *CD19* mutagenesis dataset, we comprehensively characterised the full set of splice isoforms generated in response to thousands of sequence mutations. In particular, we find that cryptic splice site activation and thus alternative 3’ and 5’ splice site usage are common modes of alternative splicing. Intriguingly, such events do not require extensive sequence remodelling, but can often be triggered by single point mutations, as indicated by strong associations between putative cryptic isoforms and certain nucleotide substitutions. This suggests, in accordance with previous reports [35], that neighbouring splice sites frequently compete for spliceosome assembly, especially if the canonical splice site is comparably weak. While this finding shows the enormous isoform complexity that can arise already from such a simple exon configuration, it raises the question of how protein function can be robustly maintained, since most cryptic *CD19* splicing isoforms likely encode non-functional proteins.

Unlike previous mutagenesis screens, which mainly focused on exonic sequence mutations, the present *CD19* dataset characterises the complete set of intronic and exonic mutations in a 1,200 nt sequence stretch. The complete characterisation of *CD19* exons 1-3 required the use of long-read sequencing technology. Given that introns in human protein-coding genes on average span ~8.1 kb (GENCODE v31), the long-read sequencing methodology described in this work opens the approach for broad applications. For *CD19*, we find that strong mutation effects are mainly centred around canonical and cryptic splice sites, whereas mutation effects seem to be dispersed for highly regulated exons such as *MSTR1* exon 11 [18]. This suggests that (near-)constitutive exons like *CD19* exon 2 may exhibit stronger and redundant splicing enhancers and that their inclusion is therefore less sensitive to individual point mutations [19]. More generally, constitutive exons may require more specific perturbations and as we show here, do not respond with only exon skipping, but tend to employ alternative splice site usage and intron retention, both of which are clinically relevant in the case of *CD19* splicing and CART-19 therapy resistance.

Our retrospective analyses of clinical B-ALL samples implicate unproductive *CD19* splice isoforms in the development of CART-19 therapy resistance. Using minigene assays, we directly show that *CD19* mutations that are observed in relapsed patients lead to exon 2 skipping, intron 2 retention or an additional isoform that uses an alternative 3’ splice site in exon 2. Furthermore, based on our mutational scan, we report ~200 additional point mutations significantly affecting these and other therapy-relevant isoforms. Taken together, our results strongly suggest that *CD19* mutations contribute to CART-19 therapy resistance by inducing splicing changes and likely do so by changing RBP binding sites in the *CD19* pre-mRNA. The detection of such mutations in longitudinal samples may provide predictive biomarkers for therapy response in the future.

At the same time, alterations in the expression of *trans*-acting RBPs can induce aberrant *CD19* splicing, explaining the presence of CD19-negative relapses in samples with a low allelic frequency of mutations or without mutations in the *CD19* locus. Mutations in splicing factors such as SRSF2, SF3B1 and U2AF1 are common in myelodysplastic syndrome/acute myelogenous leukaemia [36] and chronic lymphocytic leukaemia [37], and are associated with aberrant splicing. In B-ALL, mutations in splicing factors are not common, but previous work suggests that several splicing factors are deregulated [38]. In the context of *CD19*, we confirm that SRSF3 deregulation induces exon 2 skipping [8] and identify several other RBPs that promote CD19 protein isoforms invisible to the immunotherapeutic agent, including PTBP1, PCBP2, SF3B4, HNRNPK, MBNL1 and HNRNPM. Several of the newly identified regulators have been found as deregulated in other cancer types and are discussed as potential targets for anti-cancer therapy [39–41]. Moreover, an upregulation of PTBP1 has been implicated in the acquired resistance of pancreatic ductal carcinoma cells to the chemotherapeutic drug gemcitabine [42]. In the context of lymphocytes, PTBP1 is upregulated in B cells and required for early B cell selection [43]. It was reported, however, that treatment of leukemic cells with the targeted therapy drug imatinib, which inactivates the BCR-ABL kinase encoded by the translocated Philadelphia (Ph) chromosome, lowers PTBP1 levels [44]. In the light of our finding that *PTBP1* knockdown increases *CD19* intron 2 retention and thereby most likely reduces CD19 epitope presentation, previous treatments with imatinib may have negative impacts on subsequent responses to the CART-19 therapy in a subset of Ph+ B-ALL patients. In addition, a recent study showed that the repeat RNA *PNCTR* sequesters substantial amounts of nuclear PTBP1 in various cancers [45]. Thus, besides the regulation of protein expression, other factors like cellular availability may further impact on PTBP1 function in BALL cells under CART-19 therapy.

Currently, we cannot predict which patients with a CD19-positive B-ALL have a high risk of developing a CD19-negative relapsed disease. The pre-existence of isoforms skipping exon 2 or exons 5-6 has been previously discussed as a possible biomarker [15,16]. Our results indicate the necessity to extend the analysis to more isoforms and possibly to include the expression of splicing factors in screening approaches to identify patients at risk to relapse under CART-19 therapy. Notably, the same biomarkers might also be relevant for other malignancies arising from B-cell lineage, such as large B-cell lymphoma. Here, sequential loss of CD19 following CART-19 therapy has been described as a mechanism for relapse following immunotherapy [46], accounting for 29% of relapses in recent clinical studies [47]. Our data show that *CD19* splicing is highly complex, with already ~100 alternative isoforms concerning just exons 1-3. Of them, about 80% encode for a CD19 receptor lacking a functional CART-19 epitope and are thus expected to contribute to therapy resistance. The specific detection of alternative splicing might serve as a reliable biomarker and may provide a novel approach to monitor disease progression as already suggested in other tumour entities [48].

The contribution of aberrant splicing to CART-19 resistance may further be relevant for future combination therapies. Small-molecule splicing modulators are currently in clinical trials for myeloid neoplasms and splice site-switching antisense oligonucleotides are in development for different targets (reviewed in [11]). Our mutagenesis dataset provides a strong basis for designing and systematically evaluating splice-switching oligonucleotides for the modulation of *CD19* splicing. The combined application of these splicing modulators with immunotherapy may represent a way to limit the generation of resistance to CART therapies.

## Methods

### Cell lines

NALM-6 cells were obtained from ATCC and cultured in RPMI medium (Life Technologies) with 10% foetal bovine serum (Life Technologies) and 1% l-glutamine (Life Technologies). HEK293T cells were obtained from DSMZ and grown with the same additives as for NALM-6. All cells were kept at 37 °C in a humidified incubator containing 5% CO_2_. They were routinely tested for mycoplasma infection.

### Cloning

The *CD19* minigene was amplified from human genomic DNA (Promega) with the primers 5’-catAAGCTTgaccaccgccttcctctctg-3’ and 5’-catGAATTCNNNNNNNNNNNNNNNGGATCCttcccggcatctccccagtc-3’. pcDNA3.1 was used as the vector backbone for the *CD19* minigene plasmid. Both the backbone as well as the minigene amplicons were digested with the restriction enzymes *EcoRI* and *HindIII* (New England Biolabs). The backbone was extracted from a 1% agarose gel using QIAquick Gel Extraction Kit (Qiagen) and the minigene insert was cleaned up using QIAquick PCR Purification Kit (Qiagen). Ligation was conducted overnight at 16 °C with T4 DNA Ligase (New England Biolabs). All minigene mutations were introduced via Q5 Site-Directed Mutagenesis Kit (New England Biolabs). The nine mutations from eight patients in Orlando et al. [4] are listed in **Table S1**. All kits were used according to the manufacturers’ recommendations.

### Mutagenesis of minigene and library construction

For the random mutagenesis of the *CD19* minigene, GeneMorph II Random Mutagenesis Kit (Agilent) was used according to manufacturer’s recommendations using 500 ng *CD19* minigene for 30 cycles at 56 °C with the amplification primers 5’-catAAGCTTgaccaccgccttcctctctg-3’ and 5’-catGAATTCNNNNNNNNNNNNNNNGGATCCttcccggcatctccccagtc-3’. PCR products were purified using QIAquick Gel Extraction Kit (Qiagen), digested with *EcoR*I and *Hind*III (New England Biolabs) and then ligated into the backbone. To raise the baseline level of exon 2 inclusion in the *CD19* minigene to a similar level as in the endogenous *CD19* gene, position 748 (nucleotide 6 of intron 2) was exchanged from G to T.

### Transfection of minigene

Cells were twice washed in Dulbecco’s phosphate buffered saline (DPBS, Gibco Thermo Fisher Scientific) and then collected in R buffer with a density of 2 x 10^7^ cells/ml. For electroporation, we used 5 μg plasmid DNA (with a concentration of at least 1 μg/μl) to 2 x 10^6^ cells in R buffer for a 100 μl NEON electroporation pipette tip (Thermo Fisher Scientific) at 1600 V for 30 ms and 1 pulse. Cells were harvested 24 h later.

### Quantification of splicing isoforms using semi-quantitative RT-PCR

Semi-quantitative RT-PCR was used to quantify ratios of *CD19* mRNA isoform variants. To this end, reverse transcription was performed on 500 ng RNA with RevertAid Reverse Transcriptase (Thermo Fisher Scientific) according to the manufacturer’s recommendations. Subsequently, 1 μl of the cDNA was used as template for the RT-PCR reaction with OneTaq DNA Polymerase (New England Biolabs). PCRs were run at the following conditions: 94 °C for 30 s, 28 cycles (minigene) or 34 cycles (endogenous *CD19*) of [94 °C for 20 s, 55 °C for 30 s, 68 °C for 30 s] and final extension at 68 °C for 5 min. The primers 5’-ACCTCCTCGCCTCCTCTTCTTC-3’ and 5’-GCAACTAGAAGGCACAGTCG-3’ were used for the *CD19* minigene, and 5’-ACCTCCTCGCCTCCTCTTCTTC-3’ and 5’-CCGAAACATTCCACCGGAACAGC-3’ for the endogenous *CD19* gene. The TapeStation 2200 capillary gel electrophoresis instrument (Agilent) was used for quantification of the PCR products on D1000 tapes.

### Generation of stable and inducible shRNA knockdown cell lines

#### Production and preparation of lentivirus

Oligonucleotides with shRNA inserts against eleven RBPs (**Table S7**) were ordered as Ultramer DNA Oligos from Integrated DNA Technologies (Leuven, Belgium). All sequences were based on [49]. Oligonucleotides containing shRNA inserts were PCR-amplified with primers 5’-TCTCGAATTCTAGCCCCTTGAAGTCCGAGGCAGTAGGC-3’ and 5’-TGAACTCGAGAAGGTATATTGCTGTTGACAGTGAGCG-3’ and purified with QIAquick PCR Purification Kit (Qiagen). shRNA inserts and miRE18_LT3GEPIR_Ren714 backbone (inducible via Tet-On system) were cut with *EcoRI* and *XhoI* (New England Biolabs). Backbone was purified from agarose gel with QIAquick Gel Extraction Kit (Qiagen). The fragments were then ligated with T4 DNA Ligase (New England Biolabs) at 16 °C overnight.

Constructs were transduced into NALM-6 via HEK293T-produced lentiviruses. To this end, 10 cm dishes of HEK293T were transfected using 30 μl Lipofectamine 2000 (Thermo Fisher Scientific) with three plasmids: 4 μg shRNA-producing constructs + 2 μg psPAX2 (lentiviral packaging) + 1 μg pMD2.G (lentiviral envelope) at 72 h prior to transduction. On the first day after transfection, the medium was changed. Work with cells used for lentiviral production was conducted in the S2 laboratory.

#### Transduction of NALM-6 cells

Lentiviral production was confirmed with Lenti-X GoStix (Takara) and lentiviruses were concentrated with Lenti-X Concentrator (Takara) according to the manufacturer’s recommendations. For transduction, 1 x 10^6^ NALM-6 cells in 500 μl of medium were added to the concentrated virus. 5 μg/ml polybrene (Sigma-Aldrich) was added. The cells were centrifuged at 800 g and 32 °C for 30 min. Cells were then transferred into 6-well plates and cultivated in normal growth medium without antibiotics. Selection was started after 48 h with 0.5 μg/ml puromycin (Thermo Fisher Scientific). Antibiotic medium was exchanged every 2 to 3 days. As soon as cells were not dying under selection anymore and the population was stable, induction experiments were started. After transduction, cells remained in the S2 laboratory for at least 6 weeks. Then, Lenti-X GoStix was used to check for any remaining lentivirus.

#### Induction of stable shRNA-expressing NALM-6 cells

Controlled by the Tet-responsive *TRE3G* promoter, the expression of shRNA was induced by addition of doxycycline (Thermo Fisher Scientific). To this end, 2 x 10^6^ NALM-6 cells were seeded into a 6-well plate in 2 ml medium containing 0.5 μg/ml puromycin and induced with 0.5 μg/ml doxycycline, diluted in RPMI 1640 medium (Thermo Fisher Scientific). Induction was conducted at 37 °C and 5% CO_2_ and cells were harvested after 48 h. During induction, the shRNA expression system is coupled to the production of eGFP, which was examined by fluorescence microscopy before harvesting.

### Quantitative real-time PCR (qPCR)

RNA was extracted from the induced harvested cells using the RNeasy Plus Mini Kit (Qiagen). This RNA was used for qPCR to validate the RBP knockdown as well as for semi-quantitative RT-PCR experiments to check the splicing pattern of endogenous *CD19*. The qPCR was conducted using the Luminaris HiGreen qPCR Master Mix, low ROX (Thermo Fisher Scientific) according to the manufacturer’s recommendations. Oligonucleotide sequences of all qPCR primers are given in **Table S8**.

#### Targeted DNA sequencing

DNA-seq of the minigene library was performed on the PacBio SMRT sequencing platform at MPI-CBG Dresden. For this purpose, the minigene plasmid library was digested with *EcoR*I and *Hind*III (New England Biolabs) and run on an agarose gel. The desired band at the size of 1,301 nt was cut out and purified using QIAquick Gel Extraction Kit (Qiagen). For the run on the PacBio SMRT cell, a standard library preparation was performed.

#### Targeted RNA sequencing

NALM-6 cells were electroporated with the mutated minigene library (see above). 24 h later cells were harvested and RNA was isolated via the RNeasy Mini Kit (Qiagen). 20 μg isolated RNA was poly-A-selected using Dynabeads Oligo (dT)_25_ beads (Invitrogen) according to the manufacturer’s recommendations. Reverse transcription was performed on 500 ng poly-A-selected RNA with RevertAid Reverse Transcriptase (Thermo Fisher Scientific) according to the manufacturer’s recommendations. To prevent chimeric amplicons, the RNA-seq libraries were amplified via emulsion PCR [50] using the Phusion DNA Polymerase (New England Biolabs). The following primers containing Illumina adapters were used in the PCR: 5’-CAAGCAGAAGACGGCATACGAGATCGGTCTCGGCATTCCTGCTGAACCGCTCTTCCGATCTNNNNNNNNNNGGAACCTCTAGTGGTGAAGG-3’ (fwd) 5’-AATGATACGGCGACCACCGAGATCTACACTCTTTCCCTACACGACGCTCTTCCGATCTNNNNNNNNNNCCGCCAGTGTGATGGATATC-3’ (rev) under following conditions: 98 °C for 30 s, 25 cycles of [98 °C for 10 s, 63 °C for 20 s, 72 °C for 1 min] and final extension at 72 °C for 5 min. Amplicons were purified using Agencourt AMPure XP beads (Backman Coulter). Purified products were analysed on the TapeStation 2200 capillary gel electrophoresis instrument (Agilent) and quantified using the Qubit assay (Thermo Fisher Scientific). RNA-seq was carried out on the Illumina MiSeq platform using paired-end reads of 350 nt + 250 nt length and a 10% PhiX spike-in to increase sequence complexity.

### Re-analysis of RNA-seq data from Orlando et al

We re-analysed RNA-seq data of B-ALL patients at screening and after CART-19 therapy relapse from Orlando et al. [4] to quantify intron 2 retention in *CD19*. Since raw data were not available, we obtained BAM files for the different patients deposited in the Short Read Archive (SRA) under the accession SRP141691. For 10 patients, matched data were available at screening and relapse. The data contained the aligned reads mapped to several genes from the immune system including *CD19*. Using custom scripts, we extracted the sequence of the reads, reformatted them and generated fastq files. We then mapped the fastq files to our minigene sequence using STAR (v2.6.1) [51]. We used the re-mapped reads to quantify the levels of intron 2 retention in the different samples using the R/Bioconductor package ASpli [52].

### DNA-seq barcode demultiplexing

We obtained the circular consensus sequences (CCS), stored as fastq files. Two rounds of sequencing yielded a total of 337,215 CCS. We kept only reads with a length of 150-1,150 nt. We adapted the demultiplexing procedure described in [18]. In this case, we searched for the 15-nt barcode in the last 50 nt of the read. If the barcode was not found, we searched in the last 50 nt of the reverse complementary strand. We only allowed the recovery of barcodes ranging from 14 to 16 nt, which would account for barcodes containing one nucleotide inserted or deleted. Before proceeding with the variant calling, we determined a cutoff to decide the minimal number of CCS to call variants on. Here, we kept only barcodes supported by at least 4 CCS. In total, we recovered 68.5% of all the demultiplexed barcodes which corresponded to 10,558 different minigenes, closely resembling the ~10,000 minigene clones that were used to generate the library.

### DNA-seq mapping and variant calling

We use BLASR [53] with the standard parameters to map the de-multiplexed minigene sequences to the minigene reference. We performed variant calling in the aligned BAM files using the GATK [54] HaplotypeCaller (v4.0.10) with the parameters *--kmer-size 10 --kmer-size 15 --kmer-size 25 --allow-non-unique-kmers-in-ref*. We used different k-mer sizes to improve the detection of problematic regions. Mixed barcodes, i.e., barcodes containing two classes of mutations, were removed based on the “penetrance score”, reported as allele frequency (AF) in the GATK vcf output files, such that barcodes with more than 25% variants of low penetrance (AF < 0.8) were discarded. Using this strategy, we were able to recover 100,135 mutations of high quality coming from 10,295 distinct minigenes plus an additional 194 unmutated WT minigenes with distinct barcodes. 57.4% of the mutations appeared in at least ten different minigenes.

### RNA-seq barcode demultiplexing

RNA-seq libraries were sequenced on Illumina MiSeq as 350 nt + 250 nt paired-end reads, yielding approximately 23 million reads. We controlled their quality using FastQC (v0.11.5, https://www.bioinformatics.babraham.ac.uk/projects/fastqc/) and removed bad quality ends of reads using Trimmomatic [55] (v0.36, parameters: SLIDINGWINDOW:6:10 MINLEN:0). After trimming, we filtered for read pairs with a minimal length of 305 nt (read1) and 157 nt (read2) and, as done in Braun et al. [18], we used matchLRPatterns() and trimLRPatterns() from the R/Bioconductor package Biostrings to extract the 15 nt barcode in read1 between the two flanking restriction sites (Lpattern = TGCAGAATTC, Rpattern = GGATCC) allowing one mismatch. All read pairs with barcode length between 14 and 16 nt were kept for further processing. Barcode sequences were added to the read names in the fastq file and 5’ ends of reads were trimming (read1: everything until the second anchor sequence GGATCC, read2: the first 12 nt). After identifying and trimming the barcode and other regions, we used Cutadapt [56] (v1.6, parameters: --adapter=TAGAGGTTCC --overlap=3 --error-rate=0.1 --no-indels --minimum-length=244 --pair-filter=both) to remove remaining primer sequences from read1. Lastly, the barcode information attached to the read names was used to demultiplex all read pairs into individual fastq files for each minigene.

### Isoform quantification from RNA-seq data

Only barcodes/minigenes also detected in the DNA-seq library were kept for further analysis. All minigenes with insertions or deletions of 10 or more base pairs were removed from further analysis. For better mapping results, we shortened read1 to at most 260 nt. Read pairs of each minigene were mapped to the respective minigene (including all mutations, but excluding insertions and deletions) using STAR [51] (v2.6.1b). An annotation of three isoforms (exon 2 inclusion and skipping, as well as the artefact PCR product Δex2part which lacks an internal fragment of exon 2 due to a reverse transcription artefact [57]) was provided to STAR during mapping and an --sjdbOverhang of 259 was set. When running STAR, all SAM attributes were written, up to ten mismatches were allowed, soft-clipping was prohibited on both ends of the reads and only uniquely mapping reads were kept for further analysis. BAM files were sorted and indexed using SAMtools [58] (v1.5).

Properly and consistently mapped pairs were used for isoform reconstruction using a custom Perl script. Read pairs were considered properly mapped if they mapped with the right orientation on opposite strands. Read pairs mapped consistently if they either did not overlap or in case of an overlap, agreed in their detected splice junctions. Besides, only read pairs for which both mates exceeded the constitutive exon boundaries by at least 10 nt were used for isoform reconstruction. All other pairs were removed since they did not provide any isoform information. Only minigenes covered by at least 100 read pairs usable for isoform reconstruction were kept for further analysis. For each read pair, the CIGAR strings of the two mates were used to reconstruct their splicing isoform. Regarding the artefact product Δex2part, we combined the eight possible mappings of the missing internal fragment of exon 2 which are possible due the associated 8-nt repeat sequence [57]. Only isoforms, which were supported by >=1% of the read pairs and at least two read pairs in at least one minigene, were kept for further analysis.

The analysis described above was done separately for two replicates. All isoforms occurring with a frequency of at least 5% in two or more minigene variants in either of the two replicates were kept as individual isoforms. All other detected isoforms were summarised into a category “discarded”. Isoforms with Δex2part, i.e., excluding the internal intron in exon 2, were combined with their “real” counterparts without Δex2part by merging isoforms that only differed in the exclusion of the internal fragment of exon 2. In total, this leads to a set of 101 individual isoforms.

### Estimation of single mutation effects and splicing-effective mutations

Since the majority of the minigenes in the dataset exhibit more than one mutation, with a mean of 9.6 mutations per minigene, the splicing-effective mutations cannot be read out directly from the data. We used multinomial logistic regression to infer the effects of single mutations from combined measurements. The regression is based on hypothetical minigenes containing only one mutation, and on the assumption that mutation effects (log fold-changes compared to WT) add up into combined ones at the levels splice isoform ratios [18].

For regression, we focused on the five major isoforms that are already present in the WT minigene (see main text). Therefore, minigenes exhibiting more than 5% cryptic isoforms were removed from the dataset, and for the remaining minigenes the cryptic isoforms were merged into a lumped splicing category which we termed “other”. Thus, six categorical splicing outputs (inclusion, skipping, intron2-retention, alt-exon2, alt-exon3, other) were considered in the regression model, and the probability of each these outputs to be observed was assumed to equal the measured isoform frequencies. The regression was formulated as a softmax regression problem using the LogisticRegression command from the Python package scikit-learn [59].

Given the large number of mutations per minigene in the dataset, the regression was prone to overfitting (i.e., mutations with weak effects on splicing were assigned non-zero coefficients to fit random fluctuations in the data; not shown). To avoid this problem, we employed L1 penalisation. The strength of the penalty was optimised by tenfold cross-validation, and the resulting inverse regularisation strength was C=10 for both replicates.

The goodness of the model in describing the measured combined mutation effects (minigenes) was tested by assessing the correlation between model and data in training and test datasets (**Figure S3A**). Tenfold cross-validation at the final penalisation strength showed that the method performs very well in estimating the minigene isoform frequencies of the test dataset (**Figure S3B**). In the cross-validation, the Pearson correlation coefficients between softmax predictions of combined mutation effects and measurements lie for the single isoforms between 0.68-0.95 for the first replicate and between 0.71-0.93 for the second replicate (**Figure 3B**).

The accuracy of the model-predicted single mutation effects in the softmax regression was assessed by leaving out 56 directly measured single mutation minigenes (i.e., minigenes bearing only one mutation) from the training data. Since most of these 56 mutations are not splicing-effective, we focused our analysis on the seven mutations that change the inclusion isoform level beyond two standard deviations of the WT minigene distribution: For each of the seven mutations, we performed multiple softmax fits in which the training data: (i) contained all minigenes not harbouring the mutation of interest, (ii) excluded its single mutation minigenes, and (iii) comprised varying numbers of combined mutation minigenes containing the mutation. For each mutation occurrence between 1 and 10, we used up to 7 different, randomly chosen combinations of multiple mutated minigenes including the mutation of interest. For each of these models, we generated predictions for the single mutation effect. The prediction accuracy was assessed by calculating the difference between model and direct single mutation measurements for a certain mutation occurrence. The standard deviation of the difference between model and data was used as a measure for the model error. We find that a mutation occurrence of 3 leads to an error level equal to two WT standard deviations (calculated based on inclusion levels of all WT minigenes in the first replicate). For higher mutation occurrences, the prediction accuracy does not improve further (**Figure S3C**).

The final modelling step was to identify splicing-effective mutations. For this purpose, we adopted an approach analogous to empirical *P* values, i.e., we compared predicted single mutation effects to empirical isoform frequency distributions in the WT. Isoform frequencies were measured for 195 and 194 WT minigenes in the two replicates. For each isoform and replicate, we chose the 2.5% and 97.5% quantiles of the respective empirical WT frequency distribution as cutoffs (corresponding to a two-sided 5% cutoff). A mutation was considered to have an effect on a splice isoform if, for both replicates, the frequencies predicted by the model were beyond the respective cutoffs and if the effects were in the same direction.

### Splice site characterisation

Splice site usage for a given position represents the frequency of the isoforms using a given splice site in a particular minigene divided by the sum of all isoform frequencies for the same minigene. For **Figure 4A**, we used the maximum usage of a particular splice site across all minigenes. The strength of putative splice sites along the minigene was calculated using MaxEnt scores [25] in sliding windows of 9 nt or 23 nt to evaluate the corresponding sequences as potential 5’ or 3’ splice sites, respectively. The procedure was repeated for all individual point mutations to assess their potential to create cryptic splice sites. For the calculations we used the Python implementation of MaxEnt (maxentpy, v0.0.1, https://github.com/kepbod/maxentpy). We filtered the output by keeping only windows that contained a GU or AG dinucleotide in the positions 4-5 (5’ splice site) or 19-20 (3’ splice site), respectively.

We compared the effects of single point mutations in our library to predictions by the state-of-the-art deep learning algorithm SpliceAI [24]. We ran SpliceAI (v1.3.1) with the default parameters plus masking (-M1), using GENCODE [60] (v31) annotation for the human genome version hg38 as a reference. Given that SpliceAI results are reported in terms of a probability of gain or loss of a particular splice site, we assigned the gained splice sites in our cryptic isoforms by comparison to the canonical exon 2 inclusion isoform, such that if a new splice site appears in the cryptic isoform, it is considered as “gained” with respect to the “lost” WT splice site. All splice sites in a cryptic isoform were given the same prevalence score, i.e., the prevalence score of the mutation-isoform pair. To compare the SpliceAI scores for a given splice site gain with our prevalence score (**Figure 4F**), we considered the mutations that (i) share the same gain-loss pair of positions in both assays, and (ii) are predicted by SpliceAI to gain of a new splice site (i.e., a cryptic site where score_gain > score_loss) upon a given mutation.

### RBP binding site predictions

For the prediction of RBP binding motifs, we used the web versions of the oRNAment (http://rnabiology.ircm.qc.ca/oRNAment) [27] and ATtRACT (https://attract.cnic.es/) [26] databases to query the minigene sequence for presence of RBP motifs (**Figure S5A**). From the obtained predictions, we collapsed overlapping binding sites from the same tool and RBP.

We used DeepRiPe [28] to predict the potential impact of single point mutations on RBP binding. To this end, we downloaded the trained models for PAR-CLIP and ENCODE eCLIP data on 159 RBPs available in the Github repository (https://github.com/ohlerlab/DeepRiPe). We scored each mutation (annotated with regards to the hg38 reference genome) across the individual RBP models and preserved every mutation for which the model score changed by at least 0.25 compared to the WT sequence. The scoring functions are based on the iPython notebooks provided by DeepRiPe: https://colab.research.google.com/drive/18yeqRE7KmOjfbUaLAfJ6rMBjAulYo-Uc?usp=sharing

For the definition of significant RBP binding sites, we used the following strategy. For binding sites predicted by oRNAment and ATtRACT, we first checked their overlap separately for each isoform. If a binding site overlapped in at least one position with a splicing-effective mutation with respect to this particular isoform, we defined this binding site as an isoform-specific significant binding site. All binding sites that were significant for at least one isoform were collapsed into the complete list of significant binding sites, yielding a total of 315 significant binding sites for 74 RBPs. In the case of DeepRiPe, a mutation with a delta score > 0.25 for a given RBP model was required to overlap with a splicing-effective mutation for a particular isoform (our screen) to be considered an isoform-specific significant RBP-changing mutation. In a similar manner, all isoform-specific mutations for any isoform were collapsed into a complete list of significant RBP-changing mutations, yielding a total of 222 significant mutations that affected the binding of 58 RBPs.

### iCLIP data processing

iCLIP libraries were sequenced on an Illumina NextSeq 500 sequencing machine as 92 nt single-end reads including a 6 nt sample barcode as well as 5+4 nt unique molecular identifiers (UMIs). Basic quality controls were done with FastQC (v0.11.8) (https://www.bioinformatics.babraham.ac.uk/projects/fastqc/) and reads were filtered based on sequencing qualities (Phred score) in the barcode region using the FASTX-Toolkit (v0.0.14) (http://hannonlab.cshl.edu/fastxtoolkit/) and seqtk (v1.3) (https://github.com/lh3/seqtk/). Reads were de-multiplexed based on the experimental barcode, which is found on positions 6 to 11 of the reads, using Flexbar [61] (v3.4.0). Afterwards, barcode regions and adapter sequences were trimmed from read ends using Flexbar. Here, a minimal overlap of 1 nt of read and adapter was required, UMIs were added to the read names and reads shorter than 15 nt were removed from further analysis. Downstream analysis was done as described in Chapters 3.4 and 4.1 of Busch et al. [62]. Genome assembly and annotation of GENCODE [60] v31 were used during mapping.

### Patient data analysis

RNA-seq data of 222 B-ALL patients from the Therapeutically Applicable Research To Generate Effective Treatments (TARGET) program (https://ocg.cancer.gov/programs/target) were processed from fastq files. Sequencing adapters were trimmed with TrimGalore [63] (v0.6.6), aligned to the hg38 human genome assembly with STAR [51] (v2.5.2a), and sorted and indexed with SAMtools [58] (v1.11). Splice junctions were quantified individually for each sample using MAJIQ [64] (v2.2) and ENSEMBL reference transcriptome GRCh38.94 [65]. Only splice junctions with a usage level (percent selected index, PSI) of at least 5% in any given TARGET B-ALL samples were quantified.

## Data availability

All the sequencing data is available as a SuperSeries collection in the Gene Expression Omnibus (GEO) under the accession number GSE182894. The collection consists of the PacBio DNA-seq libraries (GSE182891), the Illumina RNA-seq libraries (GSE182892) and the PTBP1 iCLIP2 libraries in NALM-6 cells (GSE182893).

Scripts used to process the files are accessible under the GitHub repository located at: https://github.com/mcortes-lopez/CD19_splicing_mutagenesis.

The results published here are in whole or part based upon data generated by the Therapeutically Applicable Research to Generate Effective Treatments (https://ocg.cancer.gov/programs/target) initiative, phs000218. The data used for this analysis are available at https://portal.gdc.cancer.gov/projects.

## Acknowledgements

The authors would like to thank the members of the participating labs for support and discussion. We gratefully acknowledge the Institute of Molecular Biology Core Facilities for their support, especially the Genomics Core Facility and the use of its NextSeq 500 (funded by the Deutsche Forschungsgemeinschaft [DFG, German Research Foundation] INST 247/870-1 FUGG) and the Bioinformatics Core Facilities. We gratefully acknowledge the PacBio SMRT sequencing platform at MPI-CBG Dresden.

## Author contributions

M.C.-L. performed most bioinformatics analyses. L.S. performed the *CD19* minigene experiments as well as the massively parallel *CD19* splicing reporter assay. L.S. and B.S performed shRNA-mediated RBP knockdown experiments and corresponding splicing assays. M.E. and S.L. designed the mathematical modelling and prevalence score approach, and M.E. performed the analyses. F.K. contributed to quantification of mutation effects. A.O., M.C.-L., L.S. and J.K. performed PTB iCLIP experiments. A.B. performed iCLIP and RNA-seq data processing as well as splice isoform quantification. M.Q.-V. and M.T.D., performed TARGET ALL data analysis under supervision of Y.B. and A.T.-T. Study was designed by M.C.-L., L.S., M.E., K.Z., S.L. and J.K. with help from C.P., J.F. and all co-authors. K.Z., S.L. and J.K. supervised most of the bioinformatics analyses, mathematical modelling, and experimental work, respectively. M.C.-L, L.S., M.E., C.P., K.Z., S.L., and J.K. wrote the manuscript with help and comments from all co-authors.

