## Supplementary Material for "High-throughput mutagenesis identifies mutations and RNA-binding proteins controlling *CD19* splicing and CART-19 therapy resistance"

[1] Institute of Molecular Biology (IMB), Ackermannweg 4, 55128 Mainz, Germany. [2] Department of Pediatric Hematology/Oncology, Center for Pediatric and Adolescent Medicine, University Medical Center of the Johannes Gutenberg University Mainz, 55131 Mainz, Germany & University Cancer Center (UCT), University Medical Center of the Johannes Gutenberg University Mainz, 55131 Mainz & German Cancer Consortium (DKTK), site Frankfurt/Mainz, Germany, German Cancer Research Center (DKFZ), 69120 Heidelberg, Germany. [3] Department of Genetics, Perelman School of Medicine at the University of Pennsylvania, Philadelphia, PA 19104, US and Department of Biochemistry and Biophysics, Perelman School of Medicine at the University of Pennsylvania, Philadelphia, PA 19104, US. [4] Division of Cancer Pathobiology, Children's Hospital of Philadelphia, Philadelphia, PA 19104, US. [5] Department of Pathology & Laboratory Medicine, Perelman School of Medicine at the University of Pennsylvania, Philadelphia, PA 19104, US. [6] Buchmann Institute for Molecular Life Sciences (BMLS) and Faculty Biological Sciences, Goethe University Frankfurt, Max-von-Laue-Str. 15, 60438 Frankfurt, Germany. [7] Department of Systems Biology and Stuttgart Research Center for Systems Biology (SRCSB), University of Stuttgart, Stuttgart, Germany.

### These authors contributed equally.

#### SUPPLEMENTARY MATERIAL

**Table of content:**

Supplementary Figures S1-6 2

Supplementary Data S1 11

Supplementary Tables S1-8 12

Supplementary References 16

#### Supplementary Figures

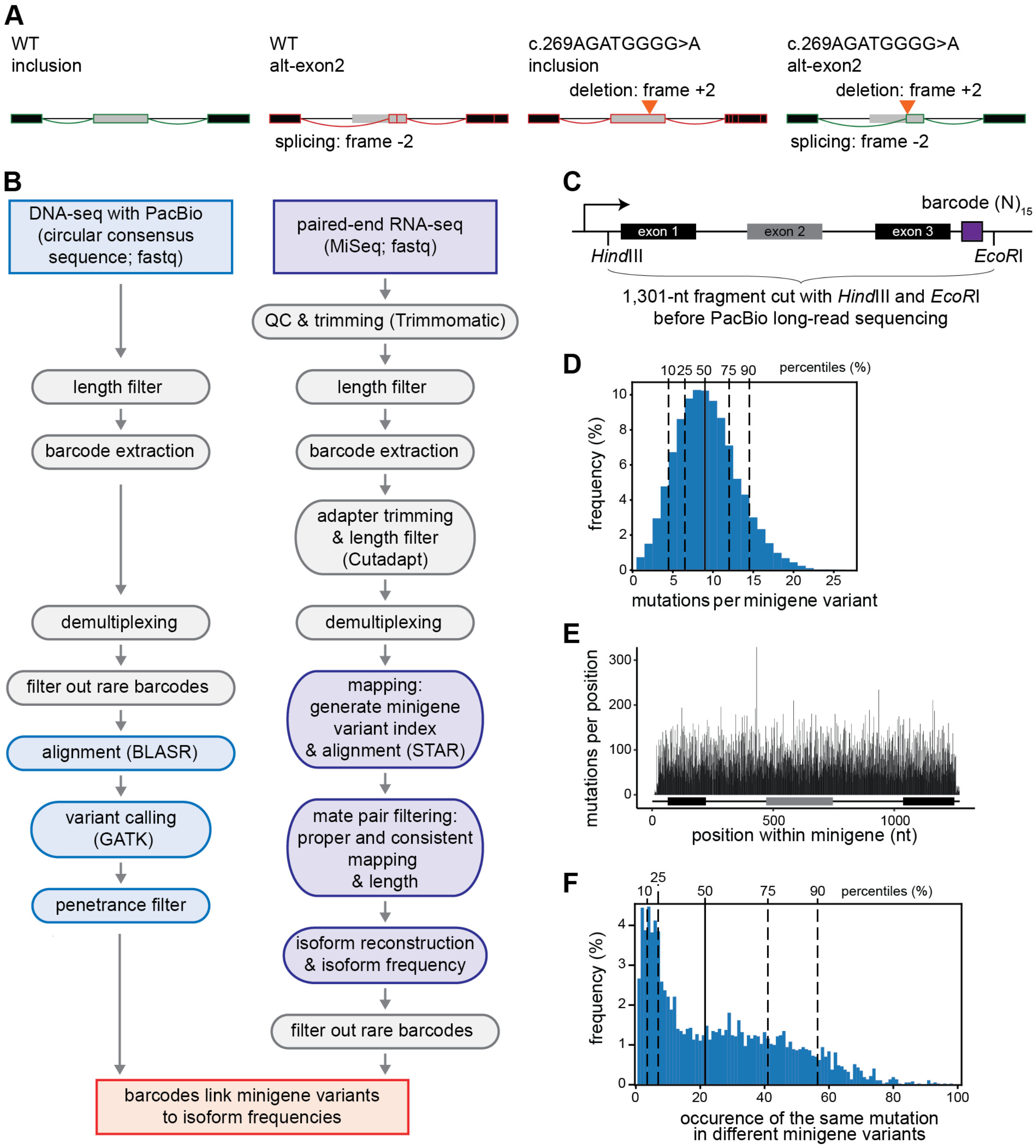


**Figure S1. Long-read sequencing identifies the introduced mutations. (A)** The deletion c.269AGATGGGG>A from patient #5 in Orlando et al. [1] introduces a frameshift (+2) that is compensated by the activation of an out-of-frame cryptic splice site (-2). Shown are the major isoforms inclusion and alt-exon2 and their coding potential in the absence (left) or presence (right) of the deletion (orange arrowhead). Schematic representation of depicts exons 1-3 (boxes) and introns (horizontal lines) with splice junctions for each isoform (arches). Colour indicates coding potential (green, coding; red, non-coding). **(B)** Analysis pipeline for the targeted DNA-seq and RNA-seq data. Left: Long-read DNA-seq data (PacBio, Pacific Bioscience) in the form of circular consensus sequences (CSS) were filtered by length (1,150-1,500 nt). 15-nt barcodes were extracted and demultiplexed, keeping only minigenes supported by at least 4 CSS. Alignment to the minigene reference was performed with BLASR [2] and variants were called using GATK HaplotypeCaller [3]. Mutations in the minigene were filtered by the “penetrance score” (allele frequency, AF), discarding all the barcodes with more than 25% variants of low penetrance (AF < 0.8). Right: Short-read RNA-seq data (Illumina) were trimmed based on quality using Trimmomatic [4] and filtered by length (305 nt for read 1, 157 nt for read 2), and adapters were trimmed using Cutadapt [5] and 15-nt barcodes were extracted and demultiplexed, keeping only minigenes supported by at least 100 read pairs. Alignment to the specific mutated version of the minigene was performed using STAR [6]. Isoform reconstruction and isoform frequency estimation was done using custom scripts (see Methods). Only minigenes with 100 or more read pairs usable for isoform reconstruction were kept. **(C)** Structure of the *CD19* minigene fragment for long-read sequencing (PacBio) to identify introduced mutations. The minigene covers exons 1-3 with the intervening introns, followed by a 15-nt barcode. The fragment for PacBio sequencing is defined by the restriction sites for *Hind*III upstream of exon 1 and *EcoR*I downstream of the barcode sequence. **(D)** 91.6% of the minigene variants carry five or more mutations. Histogram shows number of mutations per minigene for 10,295 mutated minigene variants. **(E)** 4,255 distinct mutations are spread along the *CD19* minigene, with an average of 21 mutations per position. Barplot shows the sum of mutations per position in the minigene. **(F)** 81.9% of the mutations occur in at least three minigenes, which is sufficient for a reliable estimation of single mutation effects (**Figure S3C**). Histogram shows the frequencies of the same mutations in different minigene variants.

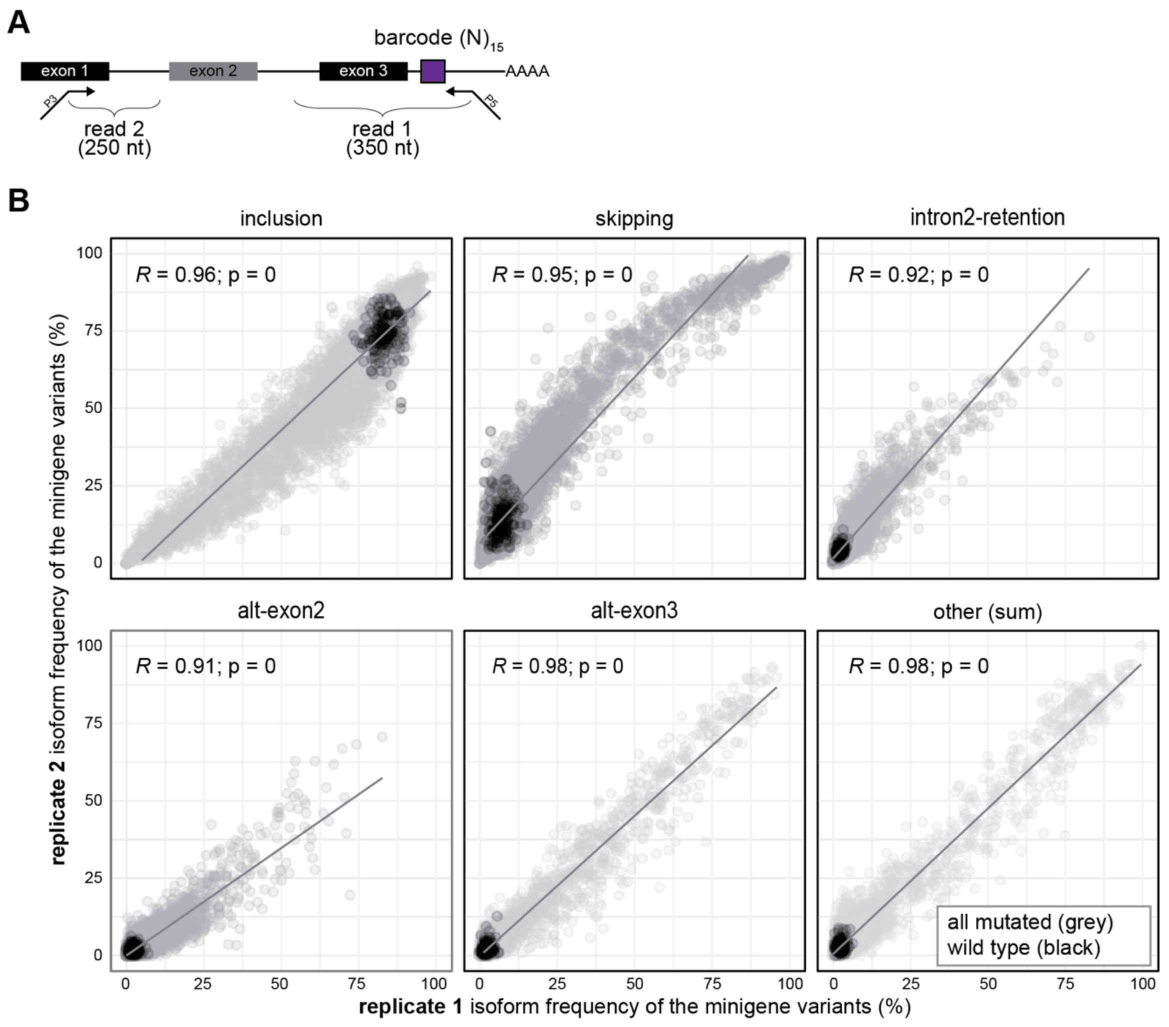


**Figure S2. Isoform measurements from targeted RNA-seq results are consistent between replicates. (A)** Description of the short-read RNA-seq strategy (Illumina) to capture the splicing products in the *CD19* minigene. Read 2 (250 nt) extends beyond exon 1, i.e. covering the exon 1/exon 2 junction, while read 1 (350 nt) includes the 15-nt barcode and extends beyond exon 3. **(B)** The isoform measurements correlate well between replicates. Scatterplots compare isoform frequencies for five major isoforms as well as the sum of 96 cryptic isoforms between replicate 1 and 2. Each dot represents a particular minigene captured in both replicates. WT and mutated minigenes appear in black and grey, respectively. Pearson correlation coefficients (*R*) and associated *P* values are given.

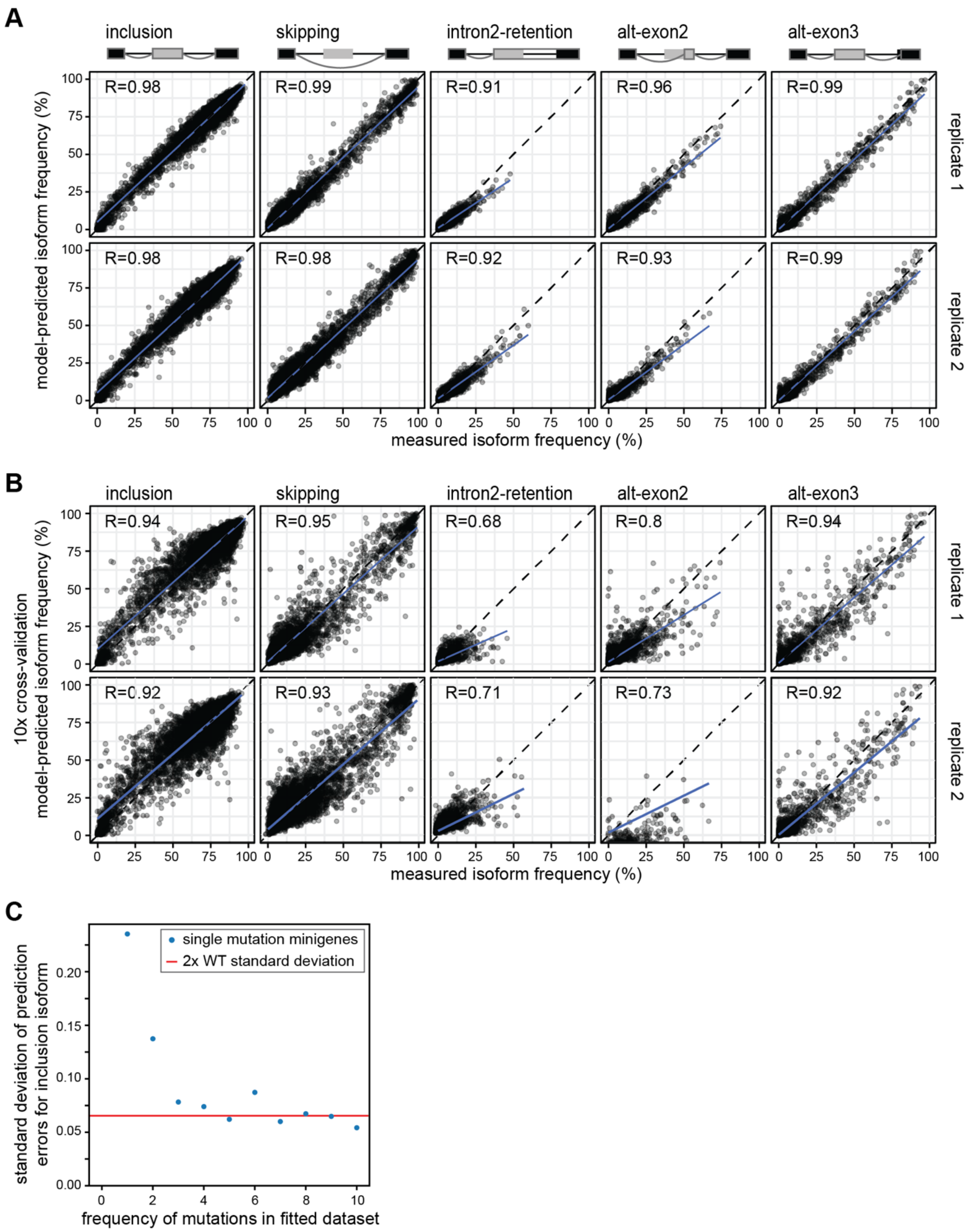


**Figure S3. The softmax regression model performs well for training and test data. (A)** Regression model fits measured combined mutation effects (i.e., minigene measurements) with high accuracy. Scatterplots show frequencies of the five major isoforms in the measurements (x-axis) against the model fit (y-axis) for two biological replicates and 9,321 minigene variants used in model training. Pearson correlation coefficients (*R*) are shown for each scatter plot. **(B)** Cross-validation confirms the predictive power of the model for minigenes not used in training. The minigene library was randomly split into ten equally sized subsets. During 10-fold cross-validation, the softmax regression model was fitted to all data excluding one subset. Scatterplots compare model-predicted splicing outcome for left-out subsets to corresponding experimental data for all major splice isoforms and are an overlay of the results of all cross-validation runs. Representation as in (A). **(C)** The model correctly infers single mutation effects. Seven single-mutation minigenes in which inclusion is significantly changed were left-out separately from softmax regression fitting and their effects were predicted based on the fit to the remaining minigene data. This procedure was repeated while additionally excluding random permutations of other minigenes containing the mutation. The standard deviation of the prediction error (y-axis) is plotted against the number of minigenes used in model training (x-axis). The inference power of the model reaches two standard deviations of the WT minigenes (horizontal line) if more than two minigenes containing the mutation are considered in model training. See Methods for details.

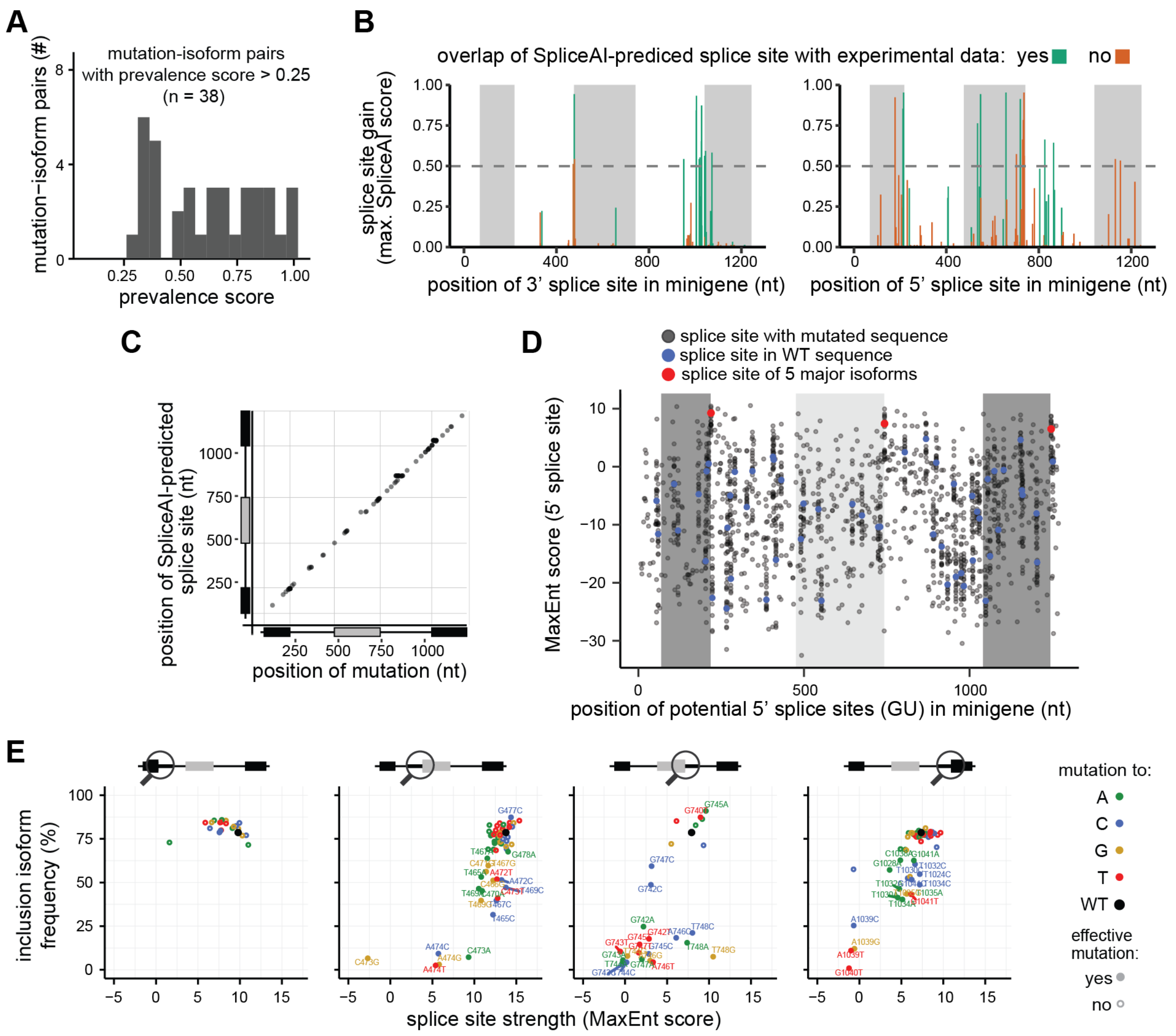


**Figure S4. Multiple mutations give rise to distinct cryptic isoforms. (A)** Multiple mutations are associated with a specific cryptic isoform. Histogram shows distribution of prevalence scores for 38 mutation-isoform pairs in which a specific mutation is associated with a distinct cryptic isoform (prevalence score > 0.25). A prevalence score of 1 indicates perfect correspondence between mutation and isoform. **(B)** SpliceAI [7] predictions for gained cryptic splice sites overlap with experimental data. Barplot shows the maximum SpliceAI score ("acceptor gain") for all the mutations that increase the probability of a given cryptic splice site to be used (38 mutations with Splice AI score [gain] > 0.5, including 15 and 23 gained 3' [left] and 5' splice sites [right]). Dotted horizontal line represents the recommended minimum threshold for a SpliceAI prediction (SpliceAI score > 0.2) [7]. Predicted gained splice sites that also appear in our experimental data are shown in green. **(C)** The mutation effects predicted by SpliceAI exclusively occur in close range, such that all SpliceAI-predicted effective mutations reside on average within 6 nt from the cryptic splice site generated. Scatterplot shows location of the gained cryptic splice sites with respect to the mutations. Only the splice site with the highest score for each mutation is considered. **(D)** The 5' splice sites of the main isoforms (red) are stronger than most other 5' splice sites in the *CD19* minigene sequence. Dotplot shows splice site strengths (MaxEnt score) [8] for putative 5' splice sites in WT (blue) and mutated (grey) minigenes in a 9-nt sliding window containing a GU dinucleotide at positions 4-5. 5' splice sites used in the five major isoforms are shown in red. **(E)** Mutation effects at 3' and 5' splice sites of *CD19* exons 2 and 3 are consistent with predicted splice site strengths. Mutations are coloured according to the changed nucleotides. Scores for WT sequence are coloured in black. Splicing-effective mutations (according to our results) are shown as filled circles and labelled, while non-effective mutations are shown as open circles.

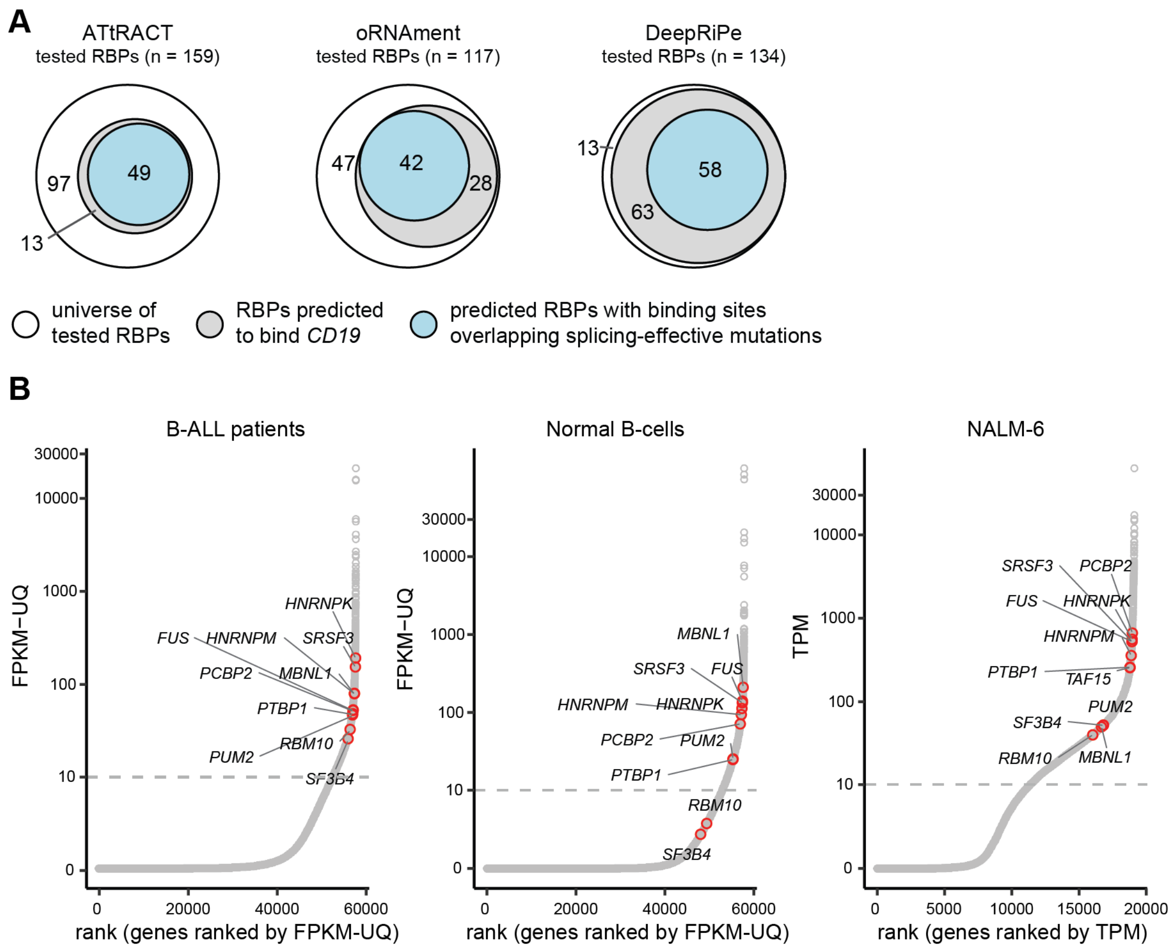


**Figure S5.** ***In silico*** **RBP binding site predictions suggest dozens of candidate regulators of** ***CD19*** **alternative splicing. (A)** *In silico* predictions of RBP binding sites were performed with ATtRACT [9] and oRNAment [10] as well as of point mutations affecting RBP binding using DeepRiPe [11]. For each prediction tool, the total number of available RBPs (white circles) is split up into those that are predicted to bind *CD19* (grey circles) and whose predicted binding sites overlap with splicing-effective mutations from our data (blue circles). Numbers refer to exclusive RBPs in each area. **(B)** Predicted RBPs were filtered based on their expression observed in B-ALL patients reported in [12]. Plot shows ranked expression values for all detected genes in samples from B-ALL patients, normal B-cells [13] and NALM-6 cells [14]. Highlighted in red are the RBP candidate genes (n = 11) tested in knockdown experiments. TPM, transcripts per million. FPKM-UQ, fragments per kilobase of transcript per million mapped reads upper quartile, a modified RNA-seq normalisation method (https://docs.gdc.cancer.gov/Encyclopedia/pages/HTSeq-FPKM-UQ/).

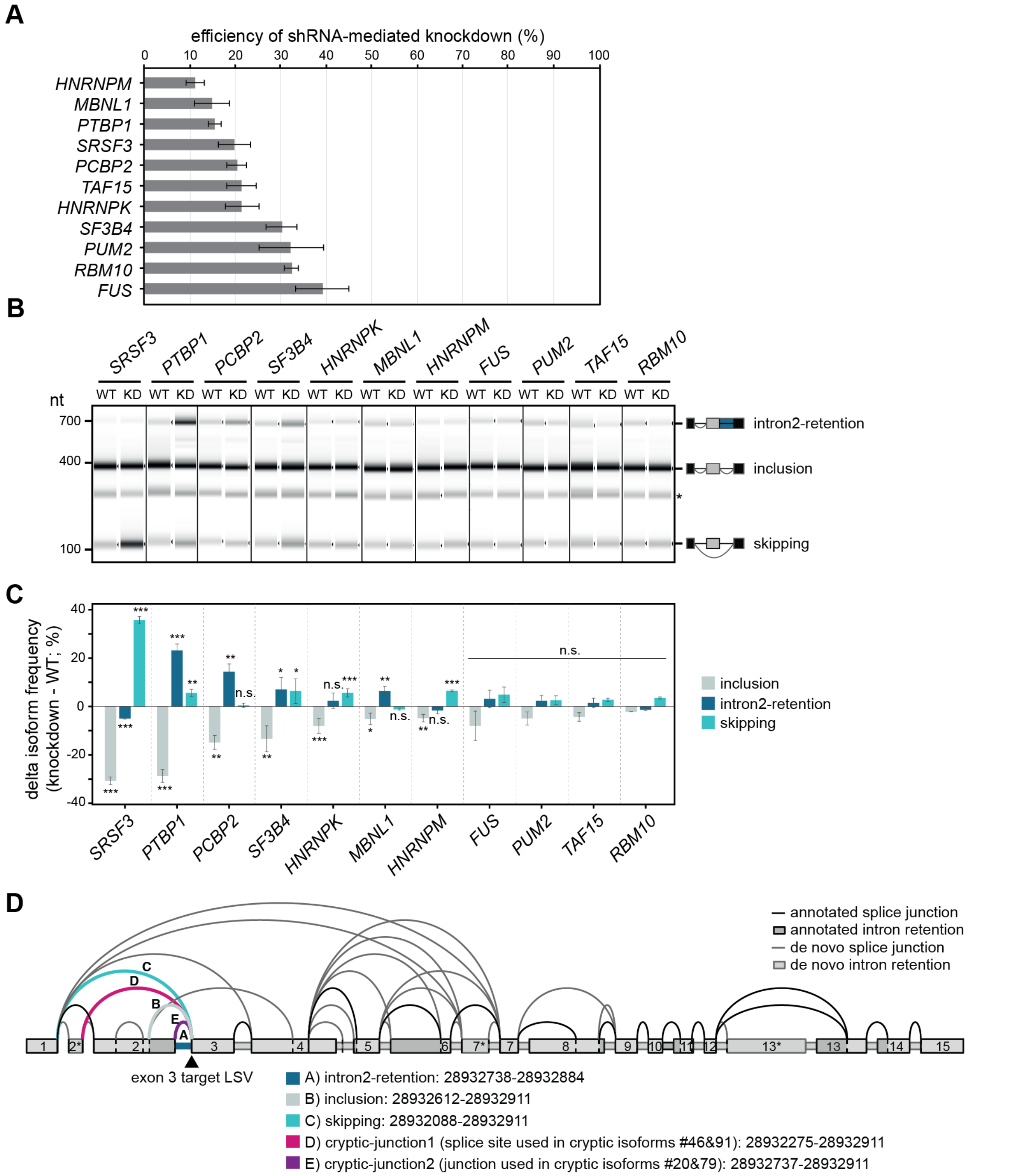


**Figure S6. Knockdown experiments show significant effects on endogenous *CD19* splicing for seven candidate RBPs. (A)** All tested RBPs are efficiently depleted upon shRNA knockdown (KD). Barplot shows mean qPCR measurements of remaining transcripts (relative to WT) for 11 candidate RBPs. Error bars indicate standard deviation of the mean (s.d.m.), n = 3 replicates. **(B, C)** Seven RBP knockdowns significantly affect *CD19* alternative splicing. Semiquantitative RT-PCR was performed to detect isoforms generated from exons 1-3 of the endogenous *CD19* gene. Gel-like representation (B), with major isoforms indicated on the right, and quantification (C), as difference in isoform frequency compared to WT, are shown. Error bars indicate s.d.m., n = 3 replicates. * *P* value < 0.05, ** *P* value < 0.01, *** *P* value < 0.01, n.s., not significant, Student's *t*-test. **(D)** *CD19* shows extensive mis-splicing in B-ALL patients. Splice junctions were quantified with MAJIQ [15] for 222 B-ALL patients from the Therapeutically Applicable Research To Generate Effective Treatments (TARGET) program (https://ocg.cancer.gov/programs/target). Splicegraph shows all splice junctions with a usage level (percent selected index, PSI) of at least 5% in any patient. Junctions and target exon of the local splicing variation (LSV) shown in **Figure 5H, I** are highlighted.

**Supplementary Data**

**Data S1. Single mutation effects on the major isoforms from the *CD19* minigene in NALM-6 cells.** For each isoform, the y-axis shows the isoform frequency (mean of two biological replicates) resulting from each individual mutation in a given position along the y-axis. Each dot represents one mutation, with colours indicating the inserted nucleotide (green, mutation to A; blue, to C; yellow, to G; red, to T). Splicing-effective mutations are shown as filled circles and non-effective mutations as open circles. Dashed lines indicate the median isoform frequency of the WT minigenes (black) ± 2 standard deviations (grey). The shown isoforms are *CD19* exon 2 inclusion, skipping, intron2-retention, alt-exon2 and alt-exon3 as well as the sum of 96 cryptic isoforms (“other”).

#### Supplementary Tables

**Table S1. Mutations from relapsed B-ALL patients reported in Orlando et al. that were tested in the *CD19* minigene splicing reporter.** Patient IDs are given as reported in Orlando et al. [1]. Note that for patient #14, two separate minigene variants were tested (#14.1 and #14.2), and that #14.2 is a combination of two adjacent mutations reported in patient #14, namely c.509A>AGTGG and c.510GCCTC>GTGGGGGAG.

| patient ID | mutation | genomic coordinate (hg38) | position in minigene | reference allele (REF) | alternative allele (ALT) |
|---|---|---|---|---|---|
| #2 | c.259G>GGGGGC | chr16:28932516 | 646 | G | GGGGGC |
| #4 | c.517TGTCTCCCACCG>T | chr16:28933072 | 1202 | TGTCTCCCACCG | T |
| #5 | c.269AGATGGGG>A | chr16:28932526 | 656 | AGATGGGG | A |
| #8 | c.265CA>C | chr16:28932522 | 652 | CA | C |
| #11 | c.264TCAACAGATGGGGGGCTTCTACCTGTGC>T | chr16:28932521 | 651 | TCAACAGATGGGGGGCTTCTACCTGTGC | T |
| #13 | c.421T>TC | chr16:28932976 | 1106 | T | TC |
| #14.1 | c.297GGGGC>G | chr16:28932554 | 684 | GGGGC | G |
| #14.2 | c.510AGCCTC>AGTGGGGGAG | chr16:28933065 | 1195 | AGCCTC | AGTGGGGGAG |
| #15 | c.271ATGGGGGGCTTCTACCTGTGCCAGCCGGGGCCC>AAGACGT | chr16:28932528 | 658 | ATGGGGGGCTTCTACCTGTGCCAGCCGGGGCCC | AAGACGT |

**Table S2. Quantification of splicing isoforms for all minigene variants in the library.** For each minigene variant, the 15-nt barcode sequence is shown together with the contained mutations, with multiple mutations separated by commas. The total number of reads per minigene variant and their distribution among the 101 isoforms are given for RNA-seq replicates 1 and 2 from NALM-6 cells. Isoform notation (219 475) indicates a splice junction that removed the region from nucleotides 219 to 475. The five major isoforms are *CD19* exon 2 inclusion (219 475)(743 1040), skipping (219 1040), intron2-retention (219 475), alt-exon2 (219 657)(743 1040) and alt-exon3 (219 475)(743 1073). In total, we detected splicing isoforms for 9,671 minigene variants in replicate 1 and for 9,372 minigene variants in replicate 2, including 9,321 minigene variants that were present in both replicates.

< provided as Excel file >

**Table S3. List of detected isoforms from the *CD19* minigene.** A total of 101 isoforms reached a relative frequency of at least 5% in at least one minigene variant, including the five major isoforms inclusion, skipping, intron2-retention, alt-exon2 and alt-exon3 (>5% in WT) as well as 96 cryptic isoforms. For each isoform, the assigned name or number is shown together with the isoform specification. Isoform notation (219 475) indicates a splice junction that removed the region from nucleotides 219 to 475. With respect to the predicted impact on the encoded CD19 protein, the number of premature stop codons (PTCs), the frame (in-frame or out-of-frame) and the resulting coding potential (coding or non-coding) are reported. With respect to an isoform's relative abundance, the average isoform frequency in the library and the maximal isoform frequency in an individual minigene are given. For the 38 cryptic isoforms that are associated with a specific mutation (prevalence score > 0.25), the respective mutations are provided together with their prevalence score and genomic coordinate (hg38). Notation G475T indicates that G in position 475 was mutated to T.

< provided as Excel file >

**Table S4. Single mutation effects predicted by the mathematical model.** Worksheet "Mutation effects" provides the model estimates of splice isoform frequencies (in %) and average delta frequency (compared to WT) in replicates (rep) 1 and 2 in response to individual mutations (single nucleotide variants, SNV; insertions or deletions, INDEL) in NALM-6 cells. Notation G475T indicates that G in position 475 was mutated to T. Individual entries are given for each affected isoform. Isoform notation (219 475) indicates a splice junction that removed the region from nucleotides 219 to 475. The five major isoforms are *CD19* exon 2 inclusion (219 475)(743 1040), skipping (219 1040), intron2-retention (219 475), alt-exon2 (219 657)(743 1040) and alt-exon3 (219 475)(743 1073). Worksheet "WT statistics" provides the mean, standard deviation (sd) and median of measured splice isoform frequencies (in %) for the five major isoforms as well as the sum of 96 cryptic isoforms ("other"). Isoform frequencies were measured for 195 and 194 WT minigenes in the two replicates.

< provided as Excel file >

**Table S5. Overlapping single nucleotide variants (SNVs) and cancer-related mutations.** Worksheet "Annotated variants" contains the SNVs (from ENSEMBL [16] v104, gnomAD [17] v3.1 and ClinVar [18] accessed 09/2021) and cancer-related variants (obtained from COSMIC [19] v94) that overlap with splicing-effective mutations and mutations with a prevalence score > 0.25 in our screen. Notation A950G indicates that A in position 950 was mutated to G. For variants present in the database dbSNP [20], the respective ID is also included. REF and ALT refer to the reference and alternative allele.

< provided as Excel file >

**Table S6. Predicted RBP binding sites in the region of the *CD19* minigene.** Worksheet "Binding sites" reports *in silico* predictions by ATtRACT [9] and oRNAment [10], providing the source tool, start and end and width (relative to the *CD19* minigene), predicted RNA-binding protein (RBP) and whether the binding site overlaps with splicing-effective mutations from our screen (see Methods). Worksheet "DeepRiPe mutations" reports all mutations predicted by DeepRiPe [11] to change RBP binding (i.e., with a delta score > 0.25), including RBP, mutation, DeepRiPe score and set as well as whether the mutation overlaps with a splicing-effective mutation from our screen and if so, for which isoform. Set refers to the DeepRiPe model that was trained for a given RBP using PAR-CLIP or ENCODE eCLIP data from HepG2 or K562 cells (see [11] for details).

< provided as Excel file >

**Table S7. Oligonucleotides used to clone the different shRNA sequence carrying vectors in this study.** Oligonucleotides were purchased from Integrated DNA Technologies.

| | |
|---|---|
| shRNA_FUS | TGCTGTTGACAGTGAGCGCACAGGATAATTCAGACAACAATAGTGAAGCCACAGATGTATTGTTGTCTGAATTATCCTGTTTGCCTACTGCCTCGGA |
| shRNA_HNRNPK | TGCTGTTGACAGTGAGCGACGAGTTGAGGCTGTTGATTCATAGTGAAGCCACAGATGTATGAATCAACAGCCTCAACTCGCTGCCTACTGCCTCGGA |
| shRNA_HNRNPM | TGCTGTTGACAGTGAGCGAAGCAGACATTCTTGAAGATAATAGTGAAGCCACAGATGTATTATCTTCAAGAATGTCTGCTCTGCCTACTGCCTCGGA |
| shRNA_MBNL1 | TGCTGTTGACAGTGAGCGCCAGCACAATGATTGACACCAATAGTGAAGCCACAGATGTATTGGTGTCAATCATTGTGCTGTTGCCTACTGCCTCGGA |
| shRNA_PCBP2 | TGCTGTTGACAGTGAGCGCTCCATCATTGAGTGTGTCAAATAGTGAAGCCACAGATGTATTTGACACACTCAATGATGGATTGCCTACTGCCTCGGA |
| shRNA_PTBP1 | TGCTGTTGACAGTGAGCGCTAGCAAGATGATACAATGGTATAGTGAAGCCACAGATGTATACCATTGTATCATCTTGCTATTGCCTACTGCCTCGGA |
| shRNA_PUM2 | TGCTGTTGACAGTGAGCGCAACATAGTTGTTGACTGTTAATAGTGAAGCCACAGATGTATTAACAGTCAACAACTATGTTATGCCTACTGCCTCGGA |
| shRNA_RBM10 | TGCTGTTGACAGTGAGCGCCGGCAAGACCATCAATGTTGATAGTGAAGCCACAGATGTATCAACATTGATGGTCTTGCCGTTGCCTACTGCCTCGGA |
| shRNA_SF3B4 | TGCTGTTGACAGTGAGCGCTGCCTTCAAGAAGGACTCCAATAGTGAAGCCACAGATGTATTGGAGTCCTTCTTGAAGGCATTGCCTACTGCCTCGGA |
| shRNA_SRSF3 | TGCTGTTGACAGTGAGCGCTAAGATGTTTTAGCTGTTCAATAGTGAAGCCACAGATGTATTGAACAGCTAAAACATCTTAATGCCTACTGCCTCGGA |
| shRNA_TAF15 | TGCTGTTGACAGTGAGCGATCAGGCTATGATCAACATCAATAGTGAAGCCACAGATGTATTGATGTTGATCATAGCCTGACTGCCTACTGCCTCGGA |

**Table S8. qPCR oligonucleotide pairs used in this study.** Oligonucleotides were purchased from Sigma-Aldrich.

| | Forward primer | Reverse primer |
|---|---|---|
| qPCR_FUS | AAGGCCTGGGTGAGAATGTT | GGCTGTCCCGTTTTCTTGTT |
| qPCR_HNRNPK | GCGAGTTGAGGCTGTTGATT | TCAGTGGAATGAGGACAGCA |
| qPCR_HNRNPM | GTCAAGGGGATGTGCTGTTG | TCCGCTCAGACTATGCTTGT |
| qPCR_MBNL1 | CGGTTTGCTCATCCTGCTGA | TTTGCACTTTTCCCGAGAGC |
| qPCR_PCBP2 | CCAGCTCTCCGGTCATCTTT | CTGGTGCAGCTTGGTCAAAT |
| qPCR_PTBP1 | CGAGATGAACACGGAGGAGG | CTGGATGTAGATGGGCTGGC |
| qPCR_PUM2 | TCAGCGTCCTCTTACTCCCA | CCAGTAGCAAGACCCTGACC |
| qPCR_RBM10 | TGTTCCCGACGTCTCTACCT | TCTCCCCATCCCAGTACAGG |
| qPCR_SF3B4 | GAACGACTTCTGGCAGCTCA | CACAGGATTGGGAGCAGAGG |
| qPCR_SRSF3 | CCCGGCTTTGCTTTTGTTGA | TTCCACTCTTACACGGCAGC |
| qPCR_TAF15 | GGTCACAGGGAGGAGGTAGA | CAGCATCTGTTCTGGGTCCA |
