## Supplementary Data S1 for "High-throughput mutagenesis identifies mutations and RNA-binding proteins controlling *CD19* splicing and CART-19 therapy resistance"

**Data S1. Single mutation effects on the major isoforms from the CD19 minigene in NALM−6 cells.**
For each isoform, the y−axis shows the isoform frequency (mean of two biological replicates) resulting from each individual mutation in a given position along the y−axis. Each dot represents one mutation, with colours indicating the inserted nucleotide (green, mutation to A; blue, to C; yellow, to G; red, to T). Splicing−effective mutations are shown as filled circles and non−effective mutations as open circles. Dashed lines indicate the median isoform frequency of the WT minigenes (black) ± 2 standard deviations (grey). The shown isoforms are CD19 exon 2 inclusion, skipping, intron2−retention, alt−exon2 and alt−exon3 as well as the sum of 96 cryptic isoforms ("other").

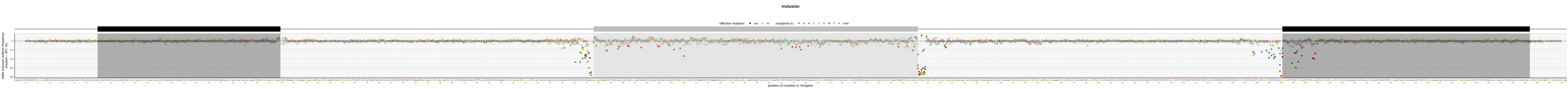

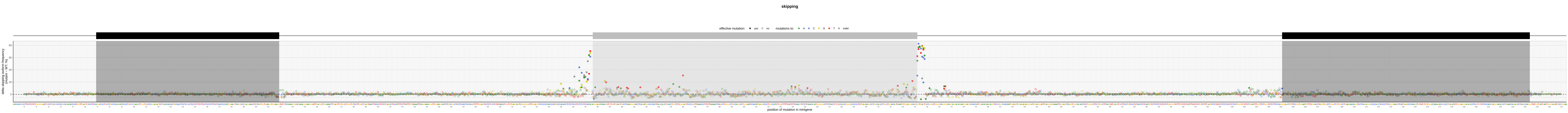

skipping
effective mutation: yes no
mutations to: A C G T indel
delta skipping isoform frequency (mutant – WT; %)
position of mutation in minigene

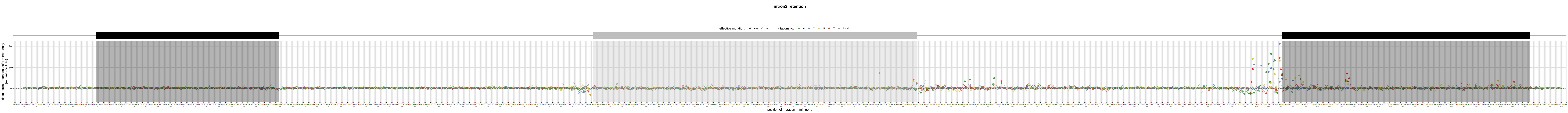
intron2 retention
effective mutation: yes no
mutations to: A C G T indel
delta intron2 retention isoform frequency (mutant - WT, %)
20
10
0
position of mutation in minigene

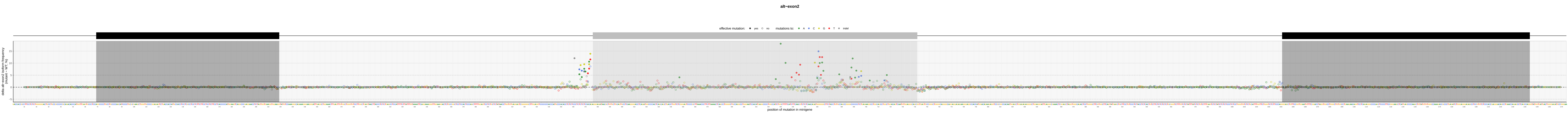
alt−exon2
effective mutation: yes no
mutations to: A C G T indel
delta alt−exon2 isoform frequency (mutant − WT; %)
15
10
5
0
−5
position of mutation in minigene

**alt−exon3**

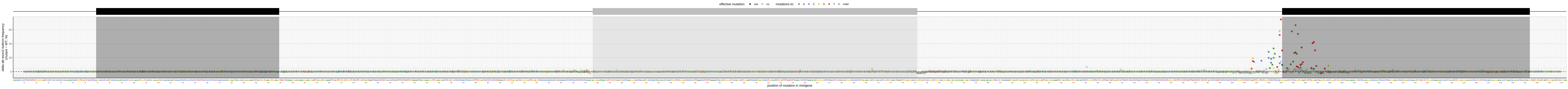
effective mutation:
yes
no
mutations to:
A
C
G
T
indel
delta alt−exon3 isoform frequency (mutant − WT; %)
0
20
40
60
position of mutation in minigene

**other**

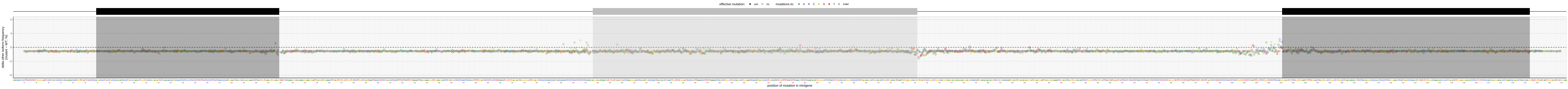

effective mutation: yes no
mutations to: A C G T indel
delta other isoform frequency (mutant – WT; %)
5
3
0
−2
−5
position of mutation in minigene
